# Visual textures are less comfortable to view than other images, consistent with efficient coding theories

**DOI:** 10.64898/2026.10.20.760531

**Authors:** Christopher DiMattina

## Abstract

Images considered uncomfortable to view often over-represent mid-range spatial frequencies (mid-SFs) relative to the average *1/f* amplitude spectrum of natural images. A proposed mechanism for this discomfort known as the Cortical Hyper-metabolism Hypothesis (CHH) posits that such images over-activate the visual cortex, which is highly sensitive to mid-SFs. We tested a novel prediction of the CHH, namely that natural image categories like textures which over-represent mid-SFs will be less comfortable to view on average than other natural images. We indeed found that texture images were less comfortable to view than various non-textures, validating this prediction of the CHH and demonstrating its potential relevance to natural image aesthetics. Passing our images through models of early visual neurons demonstrated that the mean activity of a population of simulated log-Gabor filters resembling V1 simple cells best predicted discomfort. Manipulations which selectively altered the image phase or amplitude spectra while leaving the other intact were less comfortable than the originals, suggesting that visual comfort is not a simple function of either one of these characteristics. Our results suggest that although many factors contribute to the visual comfort of natural image categories, early visual processing may play an important role, consistent with the CHH.

## INTRODUCTION

Some visual images with unnatural spatial and/or chromatic characteristics, like black and white stripes (**Fig. 1a**) or some abstract artworks, can be discomforting to view (**Fernandez & Wilkins, 2008; O’Hare & Hibbard, 2024**). In extreme cases, such images can lead some people to experience epileptiform activity or migraines (**Wilkins et al., 1984; Marcus & Soso, 1989**). One common characteristic of these images is that they often contain an excess of amplitude spectrum energy in a band of “mid-range” spatial frequencies (mid-SFs) centered around ∼3 cycles per degree (cpd) of visual angle relative to the *1/f* amplitude spectrum typical of natural images (**Wilkins et al, 1984; Fernandez & Wilkins, 2008; Wilkins & Hibbard, 2014; Wilkins, 2025; Field, 1987; Van der Schaaf & van Hateren 1996**). Given that the primate visual system is maximally sensitive to these mid-SFs (**Campbell & Robson, 1968; DeValois, Albrecht, & Thorell, 1982; Foster et al., 1985**), a mechanistic hypothesis proposed by Wilkins and colleagues (**Wilkins et al., 1984; Wilkins, 2025; Hibbard et al., 2026**) is that the over-activation of neurons in early visual cortical areas and the associated enhanced hemodynamic response is the proximate cause of the discomfort, an explanation which has been referred to as the Cortical Hyper-metabolism Hypothesis or CHH **(Wilkins & Hibbard, 2014)**. The CHH is broadly consistent with a large body of theoretical work in vision science which has suggested that the early visual system has evolved to represent natural images in an efficient manner using a minimum number of active neurons (**Barlow, 1961**; **Field 1987, 1994; Rao & Ballard, 1999; Lennie, 2003; Olshausen & Field, 2004**). Therefore, it would stand to reason that images with unnatural characteristics will not be efficiently coded by the visual system, leading to cortical over-activation and therefore visual discomfort, and possibly more serious effects like migraine or epileptic seizures (**Wilkins et al., 1979; Marcus & Soso, 1989; O’Hare & Hibbard, 2016**).

**Figure 1:**
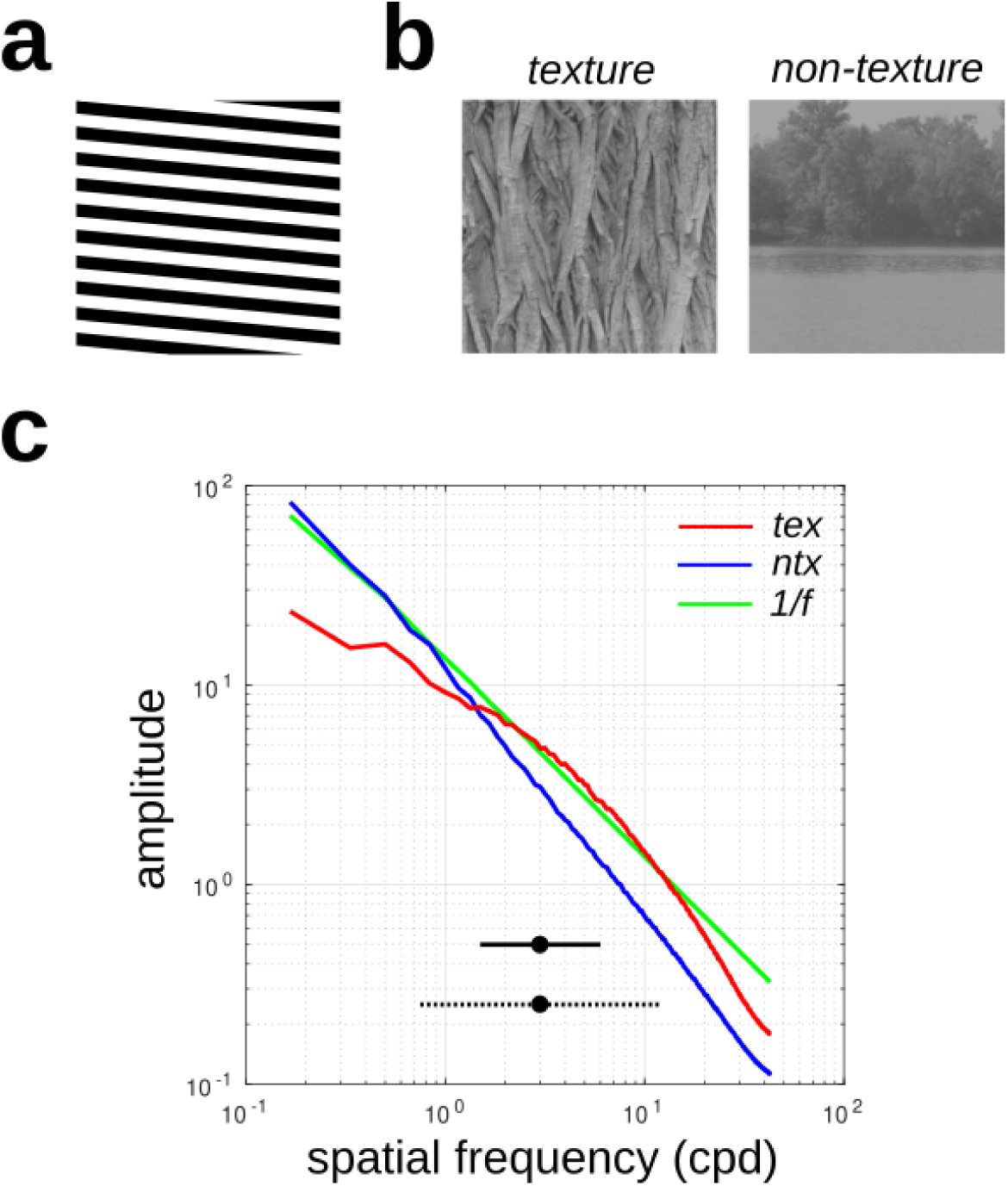
Amplitude spectra of texture stimuli and non-texture stimuli used in Experiment 1. **(a)** Example of an image containing excess Fourier energy in mid-range spatial frequencies relative to the *1/f* amplitude spectrum characteristic of most natural images. **(b)** Examples of texture (**tex**) and non-texture (**ntx**) stimuli used in **Experiment 1**. **(c)** Averaged amplitude spectra of all the *n* = 20 texture (**tex**: red) and *n* = 30 non-texture (**ntx**: blue) images, together with the amplitude spectrum of *1/f* noise (green). We find that textures over-represent spectral energy relative to *1/f* in ranges of mid-range spatial frequencies (mid-SFs) defined by 3 cpd +/− 1 octave (solid black line) and 3 cpd +/− 2 octaves (dashed black line), assuming 6 dva viewing distance. Similar results were obtained for 4, 8 dva viewing distance assumptions.

Despite its considerable support, in our opinion the predictions of the CHH have not been fully explored, with several critical questions remaining unanswered. Firstly, since different natural image categories exhibit different statistical characteristics (**Torralba & Oliva, 2003; Groen et al., 2013**), in principle this cortical hyper-metabolism hypothesis could be applied to predict and explain visual preferences for different categories of natural images. That is, given two categories of natural images, one of which is more efficiently represented in the visual cortex (less neural activity) and the other which is less efficiently represented (more neural activity), there may be a visual preference for the one which is more efficiently represented. In other words, image categories which are “easy on the eyes” are precisely those which are “easy on the brain”. Indeed, recent fMRI imaging studies have shown this to be the case (**Tang et al., 2025**), with more pleasant images being represented with less neural activity in both early visual areas (V1, V2, V4) and higher-level brain regions. In our specific context, it is known that natural visual textures over-represent energy in mid-range spatial frequencies relative to the average *1*/*f* amplitude spectrum typical of natural images (**DiMattina et al., 2024**), as we see in **Fig 1c**. Therefore, according to the CHH, textures may over-activate the cortex relative to other natural images (for instance, pictures of landscapes or objects) and be rated as less comfortable to view. This hypothesis has yet to be directly tested by comparing visual comfort ratings of natural textures to those of natural non-texture images.

Secondly, the CHH suggests that the key variable in determining cortical over-activation (and hence visual discomfort) is the amplitude spectrum of the images (**Wilkins 1984; Penacchio & Wilkins, 2015; Wilkins 2025; Hibbard et al. 2026**). However, the effects of the amplitude spectrum are ultimately mediated by the responses of populations of visual neurons. The CHH states the key variable for discomfort is the overall level of activity in the neural population, but it is possible that other population-level measures of coding efficiency, like the population kurtosis (fourth moment of the histogram of neural activities – see **Field 1987, 1994**), may better explain the effect of image amplitude spectra on visual comfort. Indeed, a previous modeling study using a simulated neural population demonstrated potential relationships between both neural population kurtosis and neural population activity and visual comfort, but it did not directly compare these predictors of discomfort (**Hibbard & O’Hare, 2015**). Therefore, it is unknown which variables describing neural population activity best predict visual comfort.

Thirdly, although the CHH is typically formulated in terms of the amplitude spectra of images, it is unclear to what extent the amplitude spectrum in and of itself is sufficient to determine visual comfort, or to what extent other factors play a role. Phase-scrambling allows one to create two images having identical amplitude spectra, but vastly different appearance due to the removal of edge structure (**Oppenheim & Lim, 1981; Pipitone & DiMattina, 2020**). If amplitude spectral characteristics are by themselves sufficient to determine visual comfort, we should expect identical comfort ratings for both original and phase-scrambled images. Likewise, given two different images, if we equalize their amplitude spectrum, we should equalize their comfort ratings. These questions have not yet been investigated in a systematic manner in the context of comparing the viewing comfort of different visual image categories.

We performed an experiment to compare the visual comfort ratings of a set of textures and non-texture images. We replicate the previously described observation of a correlation between over-representation of mid-SF energy and visual discomfort. We find that for the original images the best variable explaining visual comfort is the mean activity of a population of log-Gabor filters (**Field 1987**; **O’Hare & Hibbard, 2024**). Manipulated versions of the images which preserve the phase-spectrum and amplitude spectrum of the original were found to be less comfortable to view than the originals, demonstrating that while low-level image statistics contribute to visual comfort, they are not the primary determinant. Linear mixed effects models were shown to account for visual comfort ratings, with significant effects of image category, image manipulation, and activity of log-Gabor filters. Our results are consistent with the **CHH** in the sense that there are correlates between neural population activity (which co-varies with stimulus category) and visual comfort.

## METHODS

### Image sets and manipulations

#### Original images

We measured the visual comfort ratings of 20 natural visual textures (**tex**) and 30 natural non-textures (**ntx**). These images were all obtained from the McGill Calibrated Color Image Database (**Olmos & Kingdom, 2004**), with the 512 x 512 central region extracted from the original images (576 x 768 or 768 x 576). These images were converted to grayscale using the MATLAB function `rgb2gray.m`, and each image was set to 20% RMS contrast. Representative images from each category are shown in **Fig. 2**. The non-texture category included a variety of natural image types, including images of landscapes, plants and fruits taken at multiple scales (**Fig. 2a**). All images are freely available to the scientific community for non-commercial research purposes here: https://www.fgcu.edu/faculty/cdimattina/.

**Figure 2:**
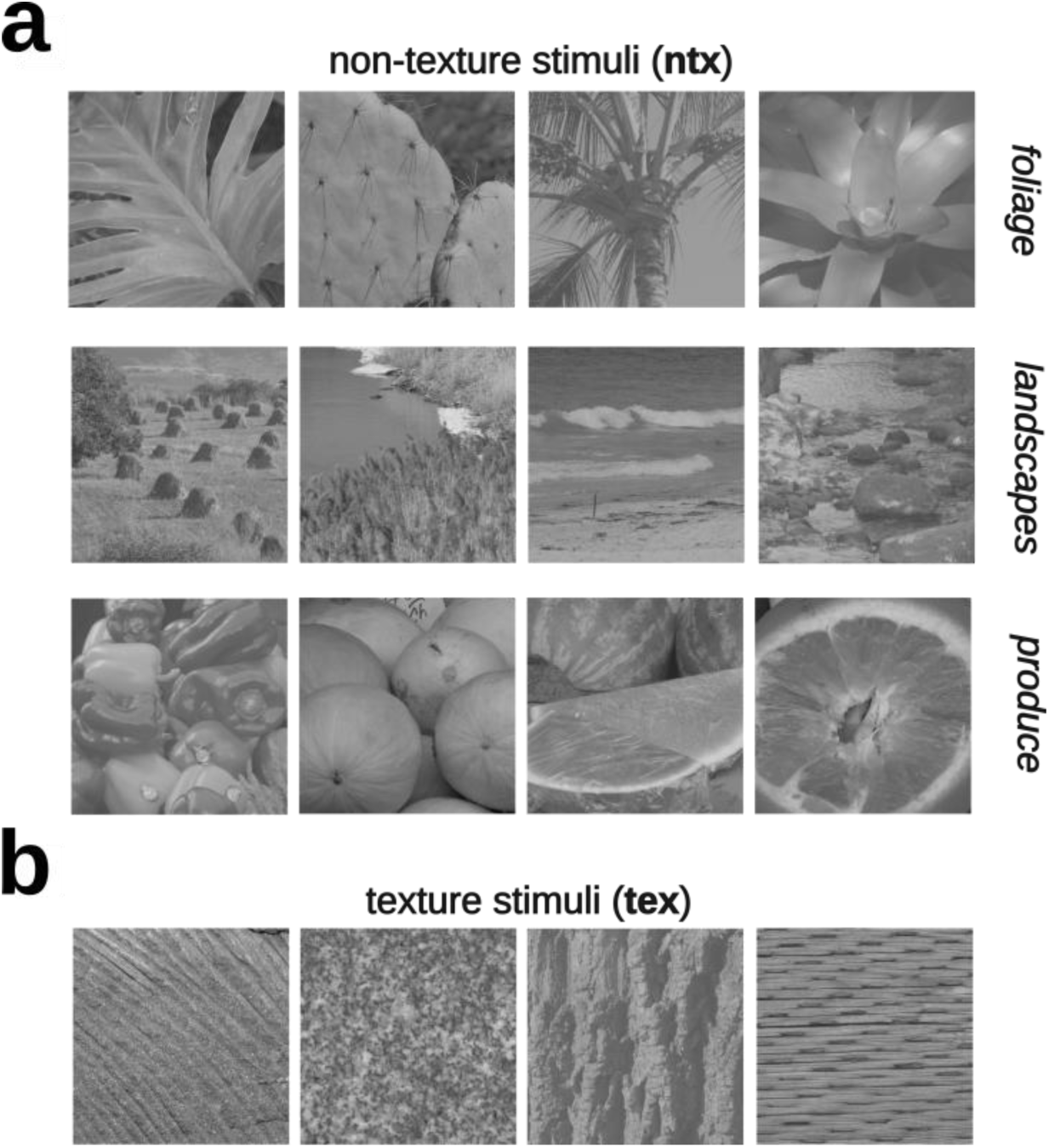
Examples of images from the (**a**) non-texture (**ntx**) and (**b**) texture (**tex**) categories.

#### Image manipulations

For both experiments, each of the original images was subject to two different manipulations to determine the relative contribution of the amplitude spectrum and phase spectrum to visual discomfort. The first manipulation was to scramble the phase of each Fourier component, while leaving the amplitude spectrum intact. This was accomplished by taking the 2D Fast Fourier Transform or FFT (using MATLAB `fft2.m`) of the original (**or**) space-domain image to transform it into the frequency domain. In the frequency domain, we break the image down into its amplitude and phase spectra. The phase spectrum is then set to that of an instantiation of Fourier transformed Gaussian white noise (which has a uniformly distributed random phase spectrum), and the resulting frequency-domain signal was then passed through the inverse 2D FFT (MATLAB function `ifft2.m`) to obtain a space-domain signal. We refer to this as the phase-scrambled (**ps**) image condition. In another manipulation, we leave the phase spectrum intact while normalizing the image amplitude spectrum to a 1/*f* shape characteristic of natural images, a condition we call **1f** or *spectrum-normalized*. Examples of all three image conditions (**or**, **ps**, **1f**) are shown in Fig. 3a (*left*).

**Figure 3:**
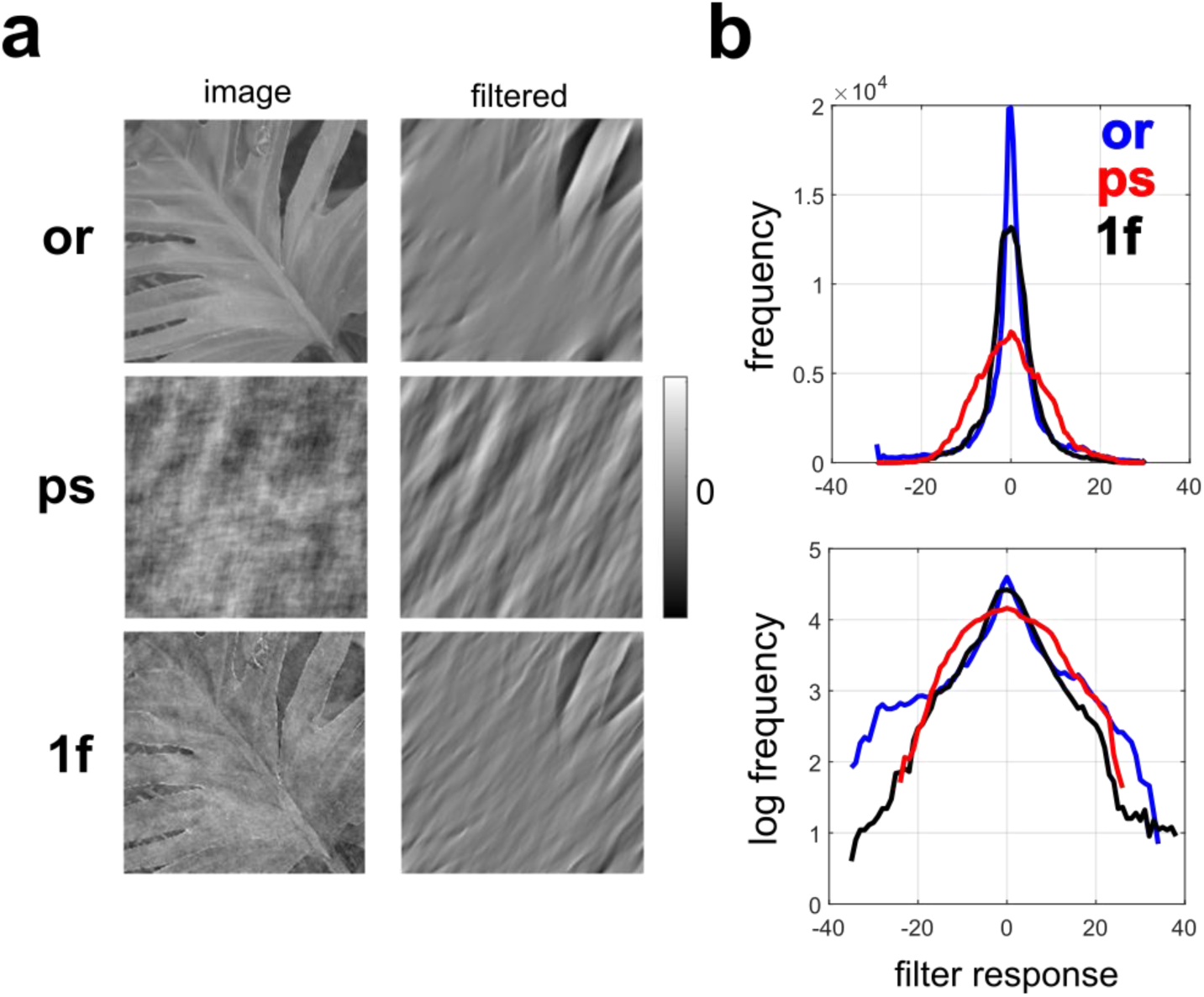
Illustration of image manipulations and responses of a Gabor filter to these images. **(a)** *Left column*: Original image (**or**, *top*) together with its phase-scrambled (**ps**, *middle*) and spectrum-normalized to *1/f* (**1f**, *bottom*) manipulations. *Right column*: The outputs of one log-Gabor filter (cell 100) from our simulated population of 500 V1 neurons when convolved with the images to the left. **(b)** Histogram of Gabor filter responses to each image shown in (a), plotted with linear (*top*) and logarithmic (*bottom*) vertical axes.

#### Amplitude spectrum analysis

According to the cortical hyper-metabolism hypothesis (**CHH**), images which over-represent mid-range spatial frequencies will over-activate early visual cortex and for this reason be uncomfortable to view (**Wilkins et al., 1984; Hibbard & Wilkins, 2014; Hibbard et al., 2026**). Therefore, following our earlier work (**DiMattina et al. 2024**), we measured the extent to which the image over-represents mid-spatial frequencies relative to the *1/f* spectrum characteristic of most natural images (**Field, 1987**). We defined a quantity called the mid-spatial frequency ratio (**mSF**) which is defined as the ratio of signal power in a specified band of frequencies to the power obtained from *1/f* noise in that same frequency band. One complication of this analysis is that the exact spatial frequency content will depend on viewing distance, and since our experiment is conducted online using Qualtrics®, participants will view the images at a variety of distances on various devices (smart phones, tablets, laptops and desktop computers). Therefore, we conduct our analysis for multiple reasonable assumptions of the size that the image subtends on the retina: 4 dva, 6 dva, 8 dva. Since our previous work found consistent results for three different definitions of mid-spatial frequencies (**DiMattina et al., 2024**), and since exact definitions of mid-spatial frequencies in the vision sciences literature vary (**Fernandez & Wilkins, 2008; Sasaki et al., 2017; Jeantet et al., 2019; Charbonneau et al., 2025**), we performed our analysis using a definition of 3 cpd +/−1 octave (1.5 to 6 cpd), as well as a broader range (3 cpd +/− 2 octaves, or 0.75 to 12 cpd). These ranges of spatial frequencies are illustrated in **Fig. 1c** (solid black line: 3 cpd +/− 1 octave, dashed black line: 3 cpd +/− 2 octaves).

### Neural population model

To test the hypothesis that images which are more comfortable to view are those which are more sparsely coded by the visual system, we applied a simple model of V1 cortical activity utilized in previous studies of visual discomfort (**O’Hare & Hibbard, 2024**). This model was comprised of log-Gabor (**Goffaux & Dakin, 2010; Field, 1987**) filters whose parameters (spatial frequency, orientation, phase) were matched to physiologically observed values (**DeValois, Albrecht & Thorell, 1982; Li, Peterson, & Freeman, 2003; Ringach, 2002**). A population of N = 500 such filters was generated and convolved with each image to obtain a distribution of V1 population responses. From this distribution, we measured two parameters: (1) The kurtosis, or equivalently, the fourth moment of the distribution, and (2) The mean absolute value of the filter response. Kurtosis (**kur-log**) measures the degree to which a distribution is peaked at zero with heavy tails, which would correspond to a sparse neural code in which most neurons exhibiting zero activity and a small number of neurons exhibiting high activity (**Field, 1987; 1994; Olshausen & Field, 2004**). The mean absolute value is a measurement of the overall activity level in the distribution (**act-log**), without regard to how it is distributed across the population.

### Behavioral experiments

To correlate neural population responses to ratings of visual discomfort, we conducted surveys using Qualtrics®. For this experiment, *n* = 132 students were recruited from Psychology classes at FGCU and completed the survey (which took less than 30 minutes) for class credit. The survey presented each of the image categories (20 texture, 30 non-texture) in all three manipulation conditions (original, phase-scrambled, *1/f*). Also presented were 10 images comprised of *1/f* noise. For each of these 160 images, observers rated the visual comfort on an 11-point Likert scale where −5 corresponds to very uncomfortable, 0 corresponds to neutral, and 5 corresponds to very comfortable. All observers provided informed consent, and the experiment was approved in advance by the FGCU Institutional Review Board (**Protocol 2022-12**), in accordance with the Declaration of Helsinki.

## RESULTS

### Analyses of image statistics and neural population encoding

#### Textures over-represent mid-SF energy

To understand the differences between texture (**tex**) and non-texture (**ntx**) images, we performed Fourier analysis on the images and characterized the responses of the neural population model (log-Gabor filters). For both definitions of mid-SF ranges (**1oct3**: 3 cpd +/− 1 octave; **2oct3**: 3 cpd +/− 2 octaves), and three assumptions about retinal image size (4, 6, 8 dva), we found significant differences between textures (**tex**, *n* = 20) and non-textures (**ntx**, *n* = 30) in the ratio of mid-SF energy relative to *1/f,* with textures consistently exhibiting higher mid-SF ratios (see **Fig. 1c**). The mid-SF (**mSF**) ratios (mean +/− SD) and independent samples t-test results for the original images (**or**) are summarized in **Table 1**.

**Table 1:** Ratio of amplitude spectral energy relative to *1/f* (mid-SF ratio: **mSF**) for textures and non-textures for three assumptions of retinal image size (4, 6, 8 dva) and two definitions (3 cpd +/−1 octaves, 3 cpd +/− 2 octaves) of mid-SF ranges (mean +/− SD). Independent samples t- tests revealed highly significant differences between non-textures (*n* = 30) and textures (*n* = 20) in all conditions.

| dva | range | non-textures | textures | *t*(48) | *p* |
|---|---|---|---|---|---|
| 4 | 1oct3 | 0.834 +/- 0.342 | 1.376 +/- 0.420 | -5.012 | < 0.001 |
|  | 2oct3 | 0.915 +/- 0.339 | 1.343 +/- 0.289 | -4.619 | < 0.001 |
| 6 | 1oct3 | 0.666 +/- 0.290 | 1.690 +/- 0.403 | -10.46 | < 0.001 |
|  | 2oct3 | 0.728 +/- 0.289 | 1.516 +/- 0.257 | -9.867 | < 0.001 |
| 8 | 1oct3 | 0.578 +/- 0.272 | 1.824 +/- 0.435 | -12.50 | < 0.001 |
|  | 2oct3 | 0.638 +/- 0.260 | 1.549 +/- 0.253 | -12.29 | < 0.001 |

Previous investigations have shown that visual textures over-represent mid-SF energy relative to the average *1/f* amplitude spectrum of natural images (**DiMattina et al., 2024**), a result which we replicate here. We see from the 95% confidence intervals of mid-SF ratios (**Table 2**) that for the texture stimuli, in all cases these intervals do not contain 1, thereby corroborating our previous results showing that textures over-represent mid-SF energy relative to *1/f*.

**Table 2:** Confidence intervals (95%) of the mean mid-SF ratios for both the non-texture stimuli (*n* = 30) and texture (*n* = 20) stimuli.

| dva | range | non-textures | textures |
|---|---|---|---|
| 4 | 1oct3 | [0.706, 0.962] | [1.180, 1.573] |
|  | 2oct3 | [0.788, 1.042] | [1.207, 1.478] |
| 6 | 1oct3 | [0.558, 0.774] | [1.501, 1.878] |
|  | 2oct3 | [0.620, 0.836] | [1.396, 1.636] |
| 8 | 1oct3 | [0.476, 0.679] | [1.621, 2.028] |
|  | 2oct3 | [0.541, 0.735] | [1.431, 1.667] |

Given the nearly perfect correlation between mid-SF ratios for the two definitions of mid-range SFs (**1oct3**, **2oct3**) at all scales analyzed (4 dva: *r* = 0.929, *p* < 0.001; 6 dva: *r* = 0.977, *p* < 0.001; 8 dva: *r* = 0.980, *p* < 0.001), all subsequent analysis is presented for the **1oct3** definition.

#### *Textures are less efficiently encoded than non-textures as measured by both population kurtosis* and activity

To understand to what extent the texture and non-texture images are coded efficiently by populations of early visual neurons, we applied the population of log-Gabor filters adapted from previous work (**Guffaux & Dakin, 2010; O’Hare & Hibbard, 2024**) to the original images (**or**) as well as their manipulations (**ps**, **1f**). One concept of efficient coding is that of *sparse coding* (**Olshausen & Field, 2004**), which is characterized by population responses exhibiting high kurtosis with a histogram of filter activities sharply peaked at zero (see **Fig. 3**), meaning that most units are inactive, with only a small number of highly active units. **Fig. 4a** (*left*) shows the population kurtosis for the log-Gabor filters for both the non-texture (**ntx**) and texture (**tex**) stimuli with 6 dva assumed image size, for all manipulation conditions (**or**: original images, **ps**: phase-scrambled, **1f**: *1/f* amplitude spectrum). Similar results were obtained for 4, 8 dva (**Supplementary Fig. S1**). We see that for the log-Gabor filters, for the original images (**or**, black circles), the kurtosis is significantly higher for the non-textures than the texture, suggesting a sparser coding of non-textures than textures. Furthermore, this difference between textures and non-textures is not simply a function of the fact that these two categories have different amplitude spectral characteristics (see previous section), as there is not a similarly large difference in population kurtosis for the phase-scrambled versions (**ps**, white squares) of the two image categories, which by definition have the same amplitude spectrum of the original images. For both textures and non-textures, we observe similarly low kurtosis for the phase-scrambled stimuli, suggesting these stimuli are not encoded efficiently for either category by this definition of coding efficiency. We see that when the phase-spectrum is preserved and the amplitude spectrum is normalized to 1/*f* for all stimuli (**1f**, white circles), that the non-texture stimuli give rise to significantly higher population kurtosis than the texture stimuli, demonstrating that the sparser representation of non-textures is largely dependent on the phase spectrum rather than the amplitude spectrum.

**Figure 4:**
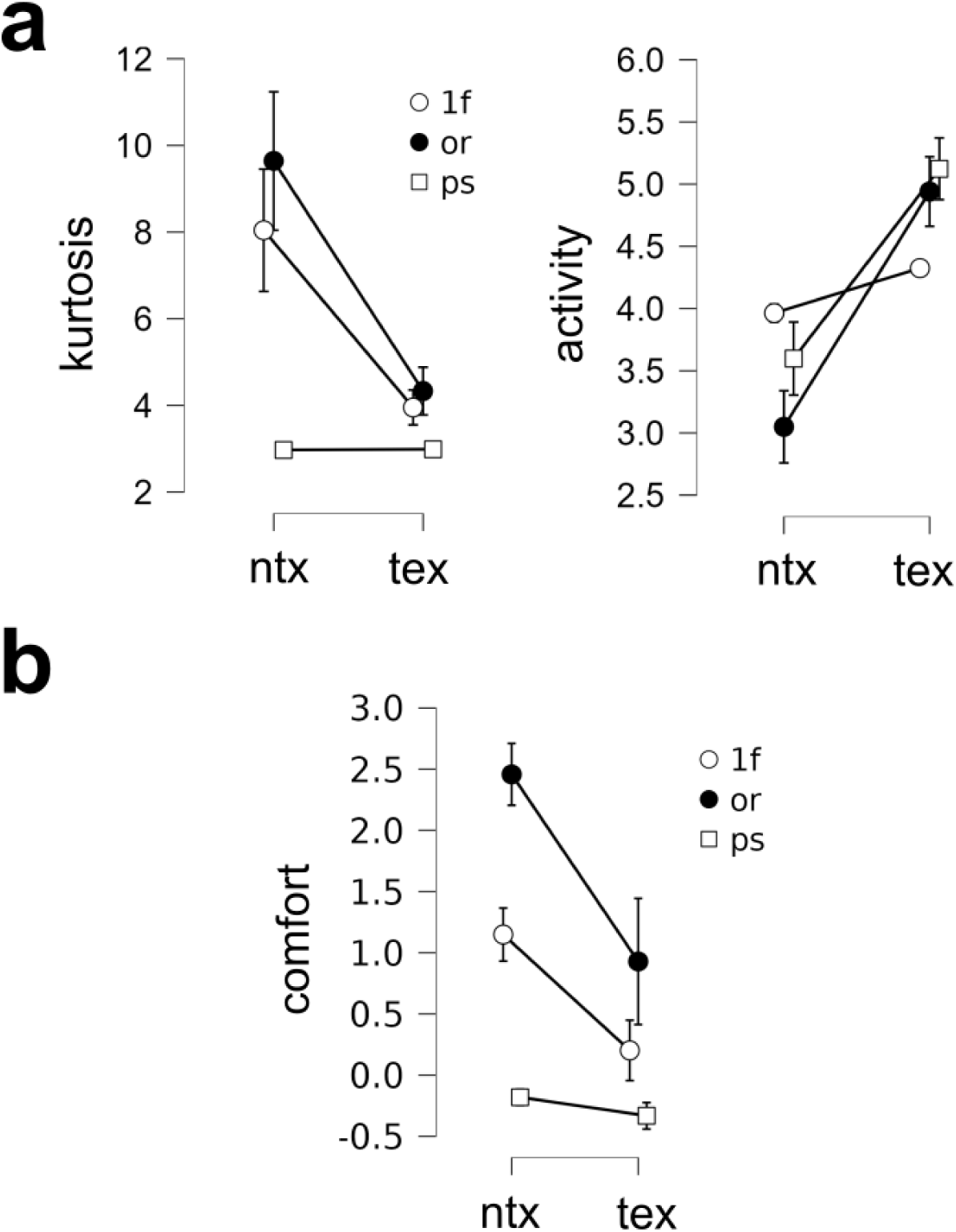
Neural coding and image viewing comfort for all image categories and manipulations. **(a)** Neural population kurtosis (*left*) and activity (*right*) for the log-Gabor population applied to all image categories (non-textures: **ntx**, textures: **tex**) and manipulations (**or**: original, **ps**: phase scrambled, **1f**: *1/f* amplitude spectrum). Error bars indicate 95% confidence intervals. Images were assumed to subtend 6 dva, and similar results were obtained for 4, 8 dva. **(b)** Average image viewing comfort for all image categories and manipulations.

Another sense in which an image can be encoded efficiently is by activating the visual cortex minimally (**Rao & Ballard 1999; Murray, Kersten & Olshausen, 2002; Hibbard et al., 2026**), and overall neural population activity can be readily measured with non-invasive methods like EEG, fMRI, and fNIRS (**O’Hare & Hibbard, 2024; Coggan et al., 2017; Wilkins 2020; Tang et al., 2025**). Therefore, we also compared the population activity elicited by the two stimulus categories (**tex**, **ntx**) for all manipulations. For the log-Gabor filters (**Fig. 4a**, *right panel*), we do see that the textures give rise to greater activity than the non-textured stimuli for the original images (**or**, black circles), and this is also true for the phase-scrambled version (**ps**, white squares), suggesting that log-Gabor population activity is modulated by image amplitude spectral characteristics. Similar results were obtained for 4, 8 dva (**Supplementary Fig. S1**). Complete ANOVA results for the plots shown in **Fig. 4a** and **Supplementary Fig. S1** are given in **Supplementary Tables S1-S6**. Interestingly, we see that the log-Gabor population seems to respond slightly more to the phase-scrambled stimuli than the original, natural stimuli (**Fig. 4a**, *right panel*). Indeed, post-hoc comparisons revealed statistically stronger responses to the phase-scrambled stimuli for all three assumptions of viewing distance, as we can see from **Supplementary Tables S2, S4, S6**.

#### Correlations between image statistics

For the original (**or**) images, we analyzed correlations between population kurtosis (**kur-log**), population activity (**act-log**), and mid-SF ratio (**mSF**). For the 6 dva viewing distance assumption, we find that **mSF** is positively correlated with **act-log** (*r* = 0.940, *p* < 0.001), negatively correlated with **kur-log** (*r* = −0.642, *p* < 0.001), and that **act-log** and **kur-log** are negatively correlated (*r* = −0.721, *p* < 0.001). Similar results were obtained for 4, 8 degrees of visual angle (**Supplementary Tables S7, S8**). We see an almost perfect correlation (*r* = 0.94) between mid-spatial frequency ratio (**mSF**) and the average activity (**act-log**) of the population of log-Gabor filters, whereas there is no significant correlation between **mSF** and the activity of the Gabor filters (**act**). This is interesting because physiological data suggests that log-Gabor filters represent a more accurate model of V1 cortical cells than Gabor filters (**Field, 1987; Hawken & Parker, 1987**). This being the case, it would suggest that images over-representing mid-spatial frequencies relative to *1/f* would over-activate the early visual cortex, a finding which has borne out in recent studies measuring cortical responses (**O’Hare et al, 2015; O’Hare, 2017; Le et al., 2017; Shi et al., 2022; Huang et al., 2011**).

### Experimental Results

#### Textures are less comfortable to view than non-textures

A 2 x 3 ANOVA revealed significant effects of image category (F_1,144_ = 74.65, *p* < 0.001, η^2^ = 0.150), manipulation (F_2,144_ = 123.30, *p* < 0.001, η^2^ = 0.497), and interaction (F_2,144_ = 15.51, *p* < 0.001, η^2^ = 0.062), on viewing comfort (**com**), as illustrated in **Fig. 4b**. For the original images (**or**, black circles), the mean viewing comfort is significantly greater (*t*(144) = 8.708, *p*_Bonf_ < 0.001, Bonferroni correction) for the non-textures (**ntx**) than for textures (**tex**).

To determine whether this difference in visual comfort was due entirely to differences in amplitude spectral content between textures and non-textures, we compared visual comfort for phase-scrambled (**ps**, white squares) versions of each image, as well as versions of each image with the amplitude spectrum fixed to the *1/f* spectrum characteristic of natural images (**1f**, white circles). As we see in **Fig. 4b**, the difference in visual comfort is not because of the amplitude spectrum, as the difference between comfort for the phase-scrambled stimuli is quite small (**ps**, white squares) and does not reach significance in our post-hoc analysis (*t*(144) = 0.864, *p*_Bonf_ = 1.00). By contrast, for the *1/f* spectrum-normalized case (**1f**, white circles) we observe a highly significant difference between the comfort of the non-texture and texture images (*t*(144) = 5.393, *p*_Bonf_ < 0.001). These results suggest that the effects of the amplitude spectrum on visual discomfort are in part dependent on the phase spectrum. We see that the **1f** images are less comfortable to view than the originals (**or**, black circles), and that the phase-scrambled images are less comfortable still (**ps**, white squares). Overall, we observe significant effect of image manipulation across categories, and interactions between category and manipulation. Complete ANOVA results for the plots shown in **Fig. 4b** are given in **Supplementary Table S9.**

Note that for the original images (**or**), although textures (**tex**) may be less comfortable than non-textures (**ntx**), the mean texture stimulus is rated as significantly more comfortable than the neutral midpoint of zero on our Likert scale (**tex**: 0.928 +/− 1.101, *t*(19) = 3.771, *p* = 0.001). That is, the original textures are not at all uncomfortable or unpleasant to view, but they are simply less pleasant than the non-textures.

#### log-Gabor filter activity best predicts mean viewing comfort for original images

For the 50 original (**or**) images of both categories: (20 **tex**, 30 **ntex**), we performed linear regression for the outcome variable of mean comfort (**com**) averaged across all *n* = 132 participants using the mid-SF ratio (**mSF**) and the neural population response characteristics (**kur-log**, **amp-log**) as predictor variables. We first reproduced the previous finding that mid-SF ratio (**mSF**) predicts mean viewing comfort (**Fernandez & Wilkins, 2008**; **Penacchio & Wilkins, 2015; Le et al., 2017**), finding that for all three assumptions of viewing size (4, 6, 8 dva) one obtains statistically significant linear model fits (4 dva: *F*_1,48_ = 6.54, *p* = 0.014, *R*^2^ = 0.120; 6 dva: *F*_1,48_ = 18.3, *p* < 0.001, *R*^2^ = 0.276; 8 dva: *F*_1,48_ = 23.4, *p* < 0.001, *R*^2^ = 0.328) with significantly negative slopes (4 dva: β = −0.866, *t(48)* = −2.56, *p* = 0.014; 6 dva: β = −0.99, *t(48)* = −4.27, *p* < 0.001; 8 dva: β = −0.93, *t(48)* = −4.84, *p* < 0.001), with models accounting for 12% to 33% of the variance in mean viewing comfort, depending on viewing distance assumption. Complete linear regression results are given in **Supplementary Tables S10-S12**.

We next sought to see whether the population of log-Gabor filters could provide a more accurate account of viewing comfort ratings. We regressed on the variables (**kur-log**, **act-log**) for all 3 assumptions of viewing distance (4, 6, 8 dva), comparing the proportions of variance accounted for by each model. In each case, we found that regressing on the kurtosis and activity of the log-Gabor population (**kur-log**, **act-log**) explained a greater degree of the variance (*R*^2^) than regressing on **mSF** (4 dva: *F*_2, 47_ = 8.160, *p* < 0.001, *R*^2^ = 0.258; 6 dva: *F*_2, 47_ = 11.716, *p* < 0.001, *R*^2^ = 0.333; 8 dva: *F*_2, 47_ = 13.977, *p* < 0.001, *R*^2^ = 0.373). Furthermore, only the variable **act-log** obtains conventional statistical significance when an individual variable is significant (4 dva: **act-log**: *p* = 0.135, **kur-log**: *p* = 0.096; 6 dva: **act-log**: *p* = 0.004, **kur-log**: *p* = 0.640; 8 dva: **act-log**: *p* < 0.001, **kur-log**: *p* = 0.906). Complete details of these regression analyses are given in **Supplementary Tables S13-S15**.

The cortical hyper-metabolism hypothesis (**CHH**) postulates that images having excessive mid-SF energy will cause excessive activation of cortical neurons (**Wilkins, 1984**), which is the cause of visual discomfort. The fact that a slightly better prediction of mean comfort (**com**) is obtained when we use **act-log** instead of **mSF** is consistent with the idea that the effects of mid-SF energy may indeed be mediated by their effects on neural activity, as suggested by the **CHH**. We tested this idea by performing a mediation analysis with independent variable **mSF**, dependent variable comfort (**com**), and mediator **act-log,** for all three assumptions of viewing distance (4, 6, 8 dva). For the original (**or**) images at all three assumptions of viewing distance (4, 6, 8 dva), we found a statistically significant indirect effect in two of the three cases (4 dva: *p* = 0.007; 6 dva: *p* = 0.042; 8 dva: *p* = 0.055) and a non-significant direct effect in all cases (4 dva, *p* = 0.217; 6 dva, *p* = 0.713; 8 dva, *p* = 0.729), consistent with the idea that the effects of **mSF** are due to the over-activation of a population of neurons in V1. Complete results of mediation analyses are given in **Supplementary Tables S16-S18**.

#### Categorical differences explain effects of act-log on mean viewing comfort

In the previous analysis, we considered all the original (**or**) images, without regard to category. However, these images were taken from two broadly defined categories: textures (**tex**) and non-textures (**ntx**). When we perform a new linear regression analysis on **act-log** which includes the variable category (**cat**) as a factor, we are able to explain more of the variance in comfort (4 dva: *R*^2^ = 0.438 vs. 0.212; 6 dva: *R*^2^ = 0.442 vs. 0.330; 8 dva: *R*^2^ = 0.447 vs. 0.373), although with the category included, the variable **act-log** no longer attains statistical significance (see **Supplementary Tables S19-S21**). This does not mean that the **act-log** variable is not important for viewing comfort – what it means is that the co-variation in **act-log** and **com** *between* categories is much greater than the co-variation *within* categories, to the point where it adds relatively little predictive power when category is included as a regressor. This is visible in the plots in **Supplementary Fig. S2**, which shows a large negative slope when images are uncategorized (*left panels*) and a smaller negative slope when image category is included as a factor (*right panels*).

#### Effects of mid-range spatial frequency for original and phase-scrambled images

We next sought to predict visual comfort using linear mixed effects modeling. Linear mixed effects modeling is a powerful technique which allows one to model relationships between fixed factors of scientific interest while controlling for sources of variability unrelated to the scientific questions, for instance inter-subject variability in comfort ratings. In this analysis, instead of predicting mean viewing comfort averaged across all participants and image exemplars, we are predicting individual viewing comfort conditioned on the participant and image exemplar. These two variables (participant, exemplar) are called the *random effects*, and the variables of interest (image category, image manipulation, neural population activity, etc…) are called *fixed effects*.

We first performed mixed-effects analysis on only the original (**or**) images. Firstly, we considered the effects of mid-range spatial frequencies when images are combined across categories, with a random effect of subject. We find that for all viewing sizes, there are significant effects of **mSF** on viewing comfort, with significant negative slopes (**Table 3**). Note that the random-effects model explains a greater proportion of the variability since it optimizes the intercept for each participant (4 dva: *R*^2^ **=** 0.344 vs. 0.120; 6 dva: *R*^2^ **=** 0.375 vs. 0.276; 8 dva: 0.385 vs. 0.328).

**Table 3:** Linear mixed-effects regression on mSF for original (or) images.

| **dva** | **R<sup>2</sup>** | **beta** | **CI-95** | **t** | **p** |
|---|---|---|---|---|---|
| 4 | 0.344 | -0.866 | [-0.977, -0.756] | -15.3 | <0.001 |
| 6 | 0.375 | -0.990 | [-1.07, -0.908] | -23.8 | < 0.001 |
| 8 | 0.385 | -0.93 | [-1.00, -0.860] | -26.2 | <0.001 |

As noted above, much of the difference in mid-range spatial frequency ratio is due to differences between the image categories (**tex**, **ntx**). Therefore, we performed mixed-effects analysis with image category as a factor and mid-spatial frequency ratio as co-variates and fit both 1-way (main effects only) and 2-way models (main effects and pairwise interactions), using the Bayes Information Criterion (**BIC**) to chose the best fitting model, penalized for complexity. As we see in **Table 4**, for both the 4 dva and 6 dva assumptions of viewing distance, we find that the preferred model (smaller BIC) has a statistically significant effect of **mSF**. as well a significant effects of category **cat**. For the 8 dva assumption of viewing distance, we obtain only a significant effect of category.

**Table 4:** Linear mixed-effects regression on cat and mSF for original images (or) with 1-way and 2-way interactions. Columns with variable names give the *p*-values for the indicated variable or interaction. Highlighted BIC values indicate the preferred model (lowest BIC).

|  | **1-way** |  |  |  | **2-way** |  |  |  |  |
|---|---|---|---|---|---|---|---|---|---|
| **dva** | **R<sup>2</sup>** | **BIC** | **cat** | **mSF** | **R<sup>2</sup>** | **BIC** | **cat** | **mSF** | **cat x mSF** |
| 4 | 0.407 | 28244 | <0.001 | 0.019 | 0.411 | **28213** | < 0.001 | 0.017 | < 0.001 |
| 6 | 0.407 | **28245** | < 0.001 | 0.033 | 0.408 | 28248 | < 0.001 | 0.073 | 0.005 |
| 8 | 0.407 | **28250** | <0.001 | 0.643 | 0.407 | 28260 | <0.001 | 0.526 | 0.463 |

Repeating the linear mixed-effects analyses performed on the original (**or**) images on the phase-scrambled (**ps**) images also revealed strong effects of mid-range spatial frequency ratio (**mSF**) on viewing comfort. Note that the models explain more of the variance in viewing comfort for the phase-scrambled images (*R*^2^ = 0.726) than the original images (*R*^2^ = 0.407), and there are stronger effects of **mSF** on viewing comfort as well. This may be because with the original images there are a wide variety of possible factors other than low-level image statistics like the amplitude spectrum which may influence viewing comfort.

**Table 5:** Linear mixed-effects regression on cat and mSF for phase-scrambled images (or) with 1-way and 2-way interactions. Columns with variable names give the *p*-values for the indicated variable or interaction. Highlighted BIC values indicate the preferred model (lowest BIC).

|  | 1-way |  |  |  | 2-way |  |  |  |  |
|---|---|---|---|---|---|---|---|---|---|
| dva | R<sup>2</sup> | BIC | cat | mSF | R<sup>2</sup> | BIC | cat | mSF | cat x mSF |
| 4 | 0.725 | **23254** | 0.651 | <0.001 | 0.725 | 23256 | 0.411 | <0.001 | 0.002 |
| 6 | 0.726 | **23227** | <0.001 | <0.001 | 0.726 | 23235 | <0.001 | <0.001 | 0.068 |
| 8 | 0.726 | **23220** | <0.001 | <0.001 | 0726 | 23220 | <0.001 | <0.001 | 0.120 |

#### Image category, manipulation, and log-Gabor filter activity predict viewing comfort

We next fit linear mixed effects models to all the data, with main effects of category (**cat**), manipulation (**man**), a **cat x man** interaction term, cluster variables of participant and image, and no co-variates. The BIC values for this model are shown in **Table 6** (*leftmost column*) for three assumptions of viewing distance. We then added single co-variates to the model to see if there was a resulting decrease in the **BIC**, which means that the addition of the co-variate improved model fits to the data while penalizing for increased model complexity. The only co-variate which we found to improve the model was the activity of the population of log-Gabor filters (**act-log**).

**Table 6:** Bayes Information Criterion (**BIC**) for linear mixed-effects regression on all the data with factors of category, manipulation and their interaction, together with a continuous co-variate added to this base mode (specified in column header). We see that the only co-variate which reduces the BIC in all cases of retinal image size assumption (4, 6, 8 dva) is the activity of the population of log-Gabor filters (**act-log**).

| dva | none | 1oct3 | kur-log | act-log |
|---|---|---|---|---|
| 4 | 81558 | 81565 | 81574 | **81553** |
| 6 | 81558 | 81562 | 81573 | **81542** |
| 8 | 81558 | 81560 | 81571 | **81536** |

We next evaluated linear mixed effects models including 1-way and 2-way interactions between **act-log** and the factors category and manipulation (**cat**, **man**). We found that for all assumptions of viewing distance, the best fitting model as measured by the BIC only included a 1-way effect of **act-log** (4 dva: 1-way = 81553, 2-way = 81580; 6 dva: 1-way: 81542, 2-way: 81563; 8 dva: 1-way: 81536, 2-way: 81554). Complete results for the 1-way linear mixed effects modeling are shown in **Supplementary Tables S22-S24**, and for the 2-way linear mixed effects models in **Supplementary Tables S25-S27**. As we see from the tables in both cases **act-log** is highly significant (*p* < 0.001). We find that the best fitting 1-way linear mixed-effects model can explain about 53% of the variance in the visual comfort ratings (4 dva: *R*^2^ = 0.531; 6 dva: *R*^2^ = 0.532; 8 dva: *R*^2^ = 0.532), with similar results obtained from the 2-way mixed effects model (4 dva: *R*^2^ = 0.532; 6 dva: *R*^2^ = 0.533; 8 dva: *R*^2^ = 0.533). **Fig. 5** shows the predictions of the best 1-way and 2-way linear mixed effects model for the 6 dva viewing size assumption. Similar results were obtained for the 4, 8 dva viewing size assumptions. We see that for the original (**or**) and spectrum-normalized (**1f**) images, that the natural images (blue) are more comfortable than the textures (yellow), whereas for the phase-scrambled images (**ps**) they are of similar comfort. Furthermore, we see that in all cases, as one increases the activity of the population of log-Gabor filters, we see a decrease in viewing comfort.

**Figure 5:**
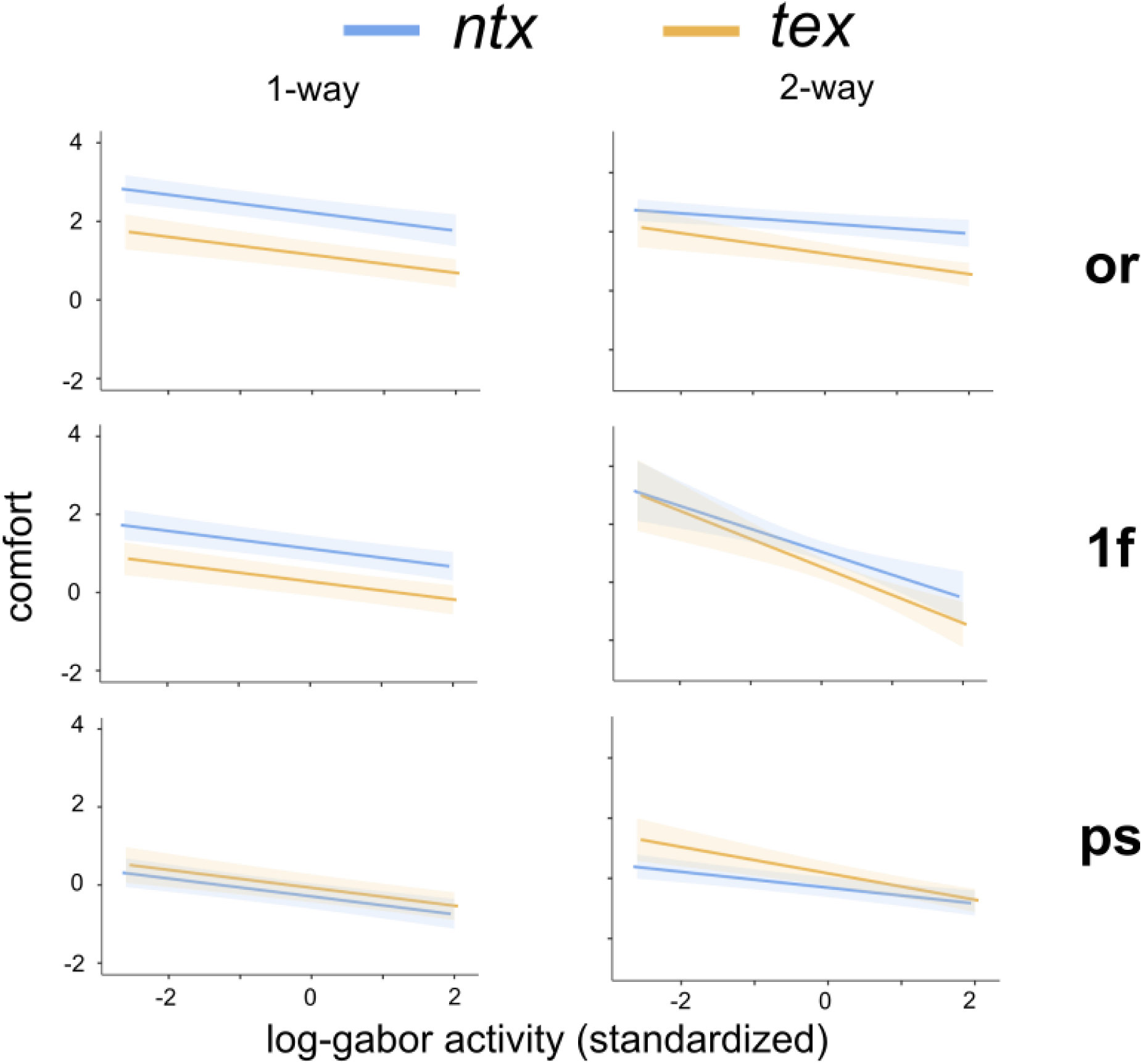
1-way and 2-way linear mixed effects model predictions for all three image manipulations and activity of the log-Gabor filters for 6 dva image size assumption.

## DISCUSSION

### Summary of findings

Visual aesthetic preferences and visual discomfort are ultimately mediated by largely unknown psychological processes and brain mechanisms (**Palmer et al., 2013; Chatterjee & Vartanian, 2016; Cardillo & Chatterjee 2026; Hibbard et al., 2026**). The cortical hyper-metabolism hypothesis (**CHH**) represents a major advance towards understanding the neural correlates of visual preferences because it relates subjective judgments of visual comfort to objectively measurable neural phenomena (**Wilkins & Hibbard, 2014; Haigh et al., 2013; Le et al., 2017; Wilkins, 2025; Hibbard et al., 2026**). Specifically, the **CHH** posits that uncomfortable images are those which over-activate early visual areas, and since the human visual system is maximally sensitive to mid-range spatial frequencies (**∼3 cpd**), images over-representing these frequencies will be uncomfortable to view.

The goal of the present study was to test one of the implicit predictions of the cortical hyper-metabolism hypothesis (**CHH**) – namely, the prediction that natural image categories which over-represent energy at mid-range spatial frequencies and over-activate early visual areas will be less comfortable to view than natural image categories which do not over-activate these areas (**Wilkins & Hibbard, 2014; Wilkins 2025**). Previous investigations revealed that visual textures over-represent mid-range spatial frequencies relative to the average 1/*f* amplitude spectrum of natural images (**DiMattina et al., 2024**), a result which we replicate here (**Tables 1-2**, **Fig. 1**). Therefore, according to the Cortical Hyper-metabolism Hypothesis (**CHH**), we expected that textures would be less comfortable to view than other image categories, and indeed we find this prediction of the **CHH** to be correct (**Fig. 4b**).

Given the strong emphasis that the **CHH** places on the spectral characteristics of images, we compared the comfort ratings of the original images (**or**) to phase-scrambled images (**ps**) with no edge structure but amplitude spectra identical to the originals, as well as to images whose amplitude spectrum was normalized to **1f**. We find that there were profound effects of these manipulations on viewing comfort. Despite having identical spectral characteristics to the original (**or**) images, the phase-scrambled (**ps**) images were rated as much less comfortable to view. This demonstrates that visual comfort is not a simple function of spectral content. Although we cannot posit a definite explanation, it is possible that the reduction in comfort comes from a mere exposure effect, which states that people find images they are familiar with or exposed to frequently to be more pleasant (**Zajonc, 1968; Cutting et al., 2006**). Since noise images are highly unnatural, it is likely that they are highly unfamiliar to most observers and hence will be rated as less pleasant. Another possibility is that these noise images may be less comfortable because they over-activate the visual cortex. Indeed, some previous work using phase-scrambled images or movies has demonstrated that they over-activate early visual areas compared to originals (**Fraedrich, et al., 2010**). We can potentially address this issue in future work using EEG or fNIRS measurements of occipital lobe activity. We also see a slight reduction in visual comfort for the **1f** images for all categories, replicating results from previous work (**Le, Cole, & Wilkins, 2015**). This again may be explained as a mere exposure effect since these images have somewhat of a “washed-out” appearance relative to typical natural images (**Fig. 3**) and are unlikely to be encountered outside of a vision science experiment.

The **CHH** posits that the variable which mediates the discomfort is the over-activation of neurons in the early visual cortex. Applying physiologically realistic models of V1 neural populations to our images to obtain predictions of neural population activity, we found a correlation between visual comfort and the activity of a population of **log-Gabor** filters (**Field, 1987**), and furthermore **log-Gabor** filter activity was shown to mediate the effects of mid-range spatial frequencies, as predicted by the **CHH**. This finding is interesting because it has been noted previously that **log-Gabor** filters provide a more physiologically realistic account of spatial frequency tuning in V1, which is bandpass on a log-frequency axis (**Hawken & Parker, 1987**). Although both kurtosis and activity exhibit correlations with viewing comfort, only activity is found in our analysis to explain discomfort. Previous work using a population of Gabor filters found that there were correlations between population kurtosis and mean activity and viewing comfort (**Hibbard & O’Hare, 2015**), but did not compare these parameters on an image-by-image basis as we do here to see which provides a better fit. Our results suggest an in important role for neural population activity, and it would be if interest to replicate this previous study using a design like the current study and see if activity of a **log-Gabor** population can better explain the variance in viewing comfort than kurtosis.

### Limitations of the current work

The current study demonstrates a correlation between the activities of a population of model neurons and visual comfort ratings. However, we must of course caution that correlation is not causation, and therefore these relationships between neural codes and visual comfort are perhaps better studied using parametric approaches with synthetic stimuli, as in previous studies of both visual discomfort (**Wilkins et al., 1984**; **O’Hare & Hibbard 2011**) and aesthetics (**e.g. Friedenberg et al., 2022**). Secondly, we rely on neural population models rather than direct measurements of visual cortex activity. It is of great interest for the near future to see if textures drive the early visual cortex more so than other categories using EEG (**O’Hare & Hibbard, 2024**; **Ogawa & Motoyoshi, 2025**), fNIRS (**Haigh et al., 2013; Le et al, 2017; Le et al. 2020**) or fMRI (**Tang et al., 2026**) measurements of stimulus-driven cortical activity. Thirdly, the present study made use of a somewhat limited set of stimuli, with 20 textures and 30 non-textures. This was done out of necessity to keep the experiment tractable in terms of time demand on participants, and to be able to explore the effects of image manipulations like phase-scrambling and amplitude spectrum normalization. Obviously, it would be of future interest to use much larger image sets from each category, perhaps making use of crowd-sourced psychophysics whereby each participant only sees a subset of stimuli. Such methods have been applied in previous studies on visual aesthetics (**Zhou et al., 2023; Tang et al., 2025**) and texture appearance (**Schmidt et al., 2025**) and could easily be applied here as well.

### Implications for future studies of texture aesthetics and visual comfort

A wide variety of factors contribute to the aesthetics of natural images, including artworks, natural scenes, and visual textures (**Palmer et al., 2013**). To the best of our knowledge, this is the first study to explicitly compare the visual comfort of texture images to non-texture images. Our finding that texture are less comfortable to view than non-textures has important implications not only for psychology and neuroscience, but fields like art, architecture, design and visual arts. Although visual discomfort is not identical to visual aesthetics, it is certainly an important piece of the puzzle. Patterns that are uncomfortable to view are rarely considered to be beautiful or chosen for decorations. Therefore, our results suggest that when choosing artwork, designers and decorators may want to chose objects or landscapes or other images with relatively slowly varying features rather than repetitive visual textures, some of which may be quite uncomfortable to view (**Valentine et al., 2026**).

Findings in the larger field of aesthetics which are not explicitly concerned with texture also have implications for texture aesthetics. For instance, it is known that sharp patterns are less comfortable to view or less pleasant than rounded things (**Bar & Neta, 2006; Vartanaian et al., 2013**). It has been suggested that this makes sense from an evolutionary perspective because sharp things like teeth or thorns can cause physical hard this may predict that texture images comprised of sharp micro-patterns may be less pleasant to view than those whose micro-patterns are more rounded, and this would be an interesting topic for future work. Other research in aesthetics which may have implications for texture aesthetics is that symmetric patterns are judged to be more pleasant (**Eisenman & Gellens, 1968; Eysesnick, 1941; Hofel & Jacobson, 2002**). Symmetry is one of several well-known Gestalt grouping cues, which describe how lower-level features are grouped into large wholes (**Palmer, 1999**). One sensible hypothesis is that textures containing various gestalt cues which enable grouping individual elements to form higher-order structure may be more comfortable to view than textures lacking such higher-level structure. This hypothesis follows from predictive coding theories of brain function which state that activity in higher brain areas predicts that in lower brain areas and suppresses the responses of neurons in lower areas whose responses are predicted (**Rao & Ballard, 1999**; **Murray, Kersten & Olshausen, 2002**). From this, it would follow that textures containing patterns which enable higher-order grouping via gestalt cues would be represented with less overall neural activity in early visual areas and may be more comfortable. Indeed, a recent groundbreaking study using fMRI demonstrated that the most comfortable images were precisely those which activated V1-V3 the least (**Tang et al., 2025**), consistent with the hypothesis that minimum energy equals visual comfort, consistent with *perceptual fluency* theories of aesthetics (**Reber, Schwartz, & Winkielman, 2004**). It would be of interest to repeat this sort of study using parametric textures whose where one can manipulate gestalt grouping cues to test this hypothesis.

Although low-level image statistics and the efficiency of the cortical representation may explain some of the aesthetic properties of textures, there are certain situations in texture perception where low-level image statistics and coding efficiency are not the full explanation. One particular category of textures which many people find unpleasant to view are those comprised of clusters of holes or bumps (**Cole & Wilkins, 2013; Cole et al., 2026**), and in a sizable minority of individual this can lead to significant distress, a condition known as *Trypophobia*. We deliberately excluded such images from our texture set here, since work from our laboratory and others suggests that Trypophobic visual discomfort is a different kind of phenomena than the visual discomfort studied here, most likely being due to an over-generalized response to images visually resembling skin disease (**Cole et al., 2026; Kupfer and Le, 2018**). Interestingly, even in studies which were not explicitly testing for Trypophobia, it was observed that patterns comprised of irregular clusters were less comfortable to view (**Ogawa, Hirata, & Motoyoshi, 2026**). These and other studies suggest that textures and materials may be able to evoke complex aesthetic and even emotional responses (**Ogawa & Motoyoshi, 2025; Ogawa et al., 2025**). Exploring the complex relationship between texture properties, visual preference, and emotional responses remains an interesting direction for future work.

## Acknowledgments

We would like to thank Sterleen Georges and Amethyst Tyler for helping with survey construction.

