## Supplementary_Material for "Visual textures are less comfortable to view than other images, consistent with efficient coding theories"

### SUPPLEMENTARY TABLE S1

**ANOVA - kurlog**

| Cases | Sum of Squares | df | Mean Square | F | p | η² |
|---|---|---|---|---|---|---|
| manip | 227.543 | 2 | 113.771 | 37.138 | < .001 | 0.236 |
| cat | 194.814 | 1 | 194.814 | 63.592 | < .001 | 0.202 |
| manip ✻ cat | 102.112 | 2 | 51.056 | 16.666 | < .001 | 0.106 |
| Residuals | 441.144 | 144 | 3.064 | | | |

*Note.* Type III Sum of Squares

**Post Hoc Comparisons - manip**

| | | Mean Difference | SE | t | $p_{bonf}$ |
|---|---|---|---|---|---|
| 1f | or | -0.608 | 0.357 | -1.703 | 0.272 |
| | ps | 2.310 | 0.357 | 6.465 | < .001 |
| or | ps | 2.918 | 0.357 | 8.168 | < .001 |

*Note.* P-value adjusted for comparing a family of 3
*Note.* Results are averaged over the levels of: cat

**Post Hoc Comparisons - manip ✻ cat**

| | | Mean Difference | SE | t | $p_{bonf}$ |
|---|---|---|---|---|---|
| 1f N | or N | -0.938 | 0.452 | -2.076 | 0.595 |
| | ps N | 3.908 | 0.452 | 8.648 | < .001 |
| | 1f T | 3.172 | 0.505 | 6.278 | < .001 |
| | or T | 2.893 | 0.505 | 5.727 | < .001 |
| | ps T | 3.883 | 0.505 | 7.686 | < .001 |
| or N | ps N | 4.846 | 0.452 | 10.724 | < .001 |
| | 1f T | 4.110 | 0.505 | 8.135 | < .001 |
| | or T | 3.832 | 0.505 | 7.583 | < .001 |
| | ps T | 4.822 | 0.505 | 9.543 | < .001 |
| ps N | 1f T | -0.736 | 0.505 | -1.457 | 1.000 |
| | or T | -1.015 | 0.505 | -2.009 | 0.697 |
| | ps T | -0.025 | 0.505 | -0.049 | 1.000 |
| 1f T | or T | -0.279 | 0.553 | -0.503 | 1.000 |
| | ps T | 0.711 | 0.553 | 1.285 | 1.000 |
| or T | ps T | 0.990 | 0.553 | 1.789 | 1.000 |

*Note.* P-value adjusted for comparing a family of 6

**Supplementary Table S1:** ANOVA for the effects of image category (non-texture: **N**, texture: **T**) and manipulation (**or**: original, **ps**: phase-scrambled, **1f**: amplitude-normalized) on the kurtosis of the log-Gabor filter population (**kurlog**) for the assumed viewing distance of 4 dva.

### SUPPLEMENTARY TABLE S2

**ANOVA - actlog**

| Cases | Sum of Squares | df | Mean Square | F | p | η² |
|---|---|---|---|---|---|---|
| manip | 2.420 | 2 | 1.210 | 3.497 | 0.033 | 0.030 |
| cat | 22.687 | 1 | 22.687 | 65.565 | < .001 | 0.285 |
| manip ✻ cat | 4.702 | 2 | 2.351 | 6.794 | 0.002 | 0.059 |
| Residuals | 49.827 | 144 | 0.346 | | | |

*Note.* Type III Sum of Squares

**Post Hoc Comparisons - manip**

| | | Mean Difference | SE | t | $p_{bonf}$ |
|---|---|---|---|---|---|
| 1f | or | 0.121 | 0.120 | 1.006 | 0.949 |
| | ps | -0.194 | 0.120 | -1.616 | 0.325 |
| or | ps | -0.315 | 0.120 | -2.621 | 0.029 |

*Note.* P-value adjusted for comparing a family of 3
*Note.* Results are averaged over the levels of: cat

**Post Hoc Comparisons - manip ✻ cat**

| | | Mean Difference | SE | t | $p_{bonf}$ |
|---|---|---|---|---|---|
| 1f N | or N | 0.561 | 0.152 | 3.694 | 0.005 |
| | ps N | 0.066 | 0.152 | 0.433 | 1.000 |
| | 1f T | -0.327 | 0.170 | -1.927 | 0.840 |
| | or T | -0.647 | 0.170 | -3.808 | 0.003 |
| | ps T | -0.781 | 0.170 | -4.599 | < .001 |
| or N | ps N | -0.495 | 0.152 | -3.261 | 0.021 |
| | 1f T | -0.888 | 0.170 | -5.231 | < .001 |
| | or T | -1.208 | 0.170 | -7.112 | < .001 |
| | ps T | -1.342 | 0.170 | -7.902 | < .001 |
| ps N | 1f T | -0.393 | 0.170 | -2.314 | 0.331 |
| | or T | -0.712 | 0.170 | -4.195 | < .001 |
| | ps T | -0.847 | 0.170 | -4.986 | < .001 |
| 1f T | or T | -0.319 | 0.186 | -1.718 | 1.000 |
| | ps T | -0.454 | 0.186 | -2.439 | 0.239 |
| or T | ps T | -0.134 | 0.186 | -0.721 | 1.000 |

*Note.* P-value adjusted for comparing a family of 6

**Supplementary Table S2:** ANOVA for the effects of image category (non-texture: **N**, texture: **T**) and manipulation (**or**: original, **ps**: phase-scrambled, **1f**: amplitude-normalized) on the activity of the log-Gabor filter population (**actlog**) for the assumed viewing distance of 4 dva.

#### SUPPLEMENTARY TABLE S3

**ANOVA - kurlog**

| Cases | Sum of Squares | df | Mean Square | F | p | $\eta^2$ |
|---|---|---|---|---|---|---|
| manip | 418.123 | 2 | 209.061 | 30.583 | < .001 | 0.215 |
| cat | 352.133 | 1 | 352.133 | 51.512 | < .001 | 0.181 |
| manip ✻ cat | 186.806 | 2 | 93.403 | 13.664 | < .001 | 0.096 |
| Residuals | 984.368 | 144 | 6.836 | | | |

*Note.* Type III Sum of Squares

**Post Hoc Comparisons - manip**

| | | Mean Difference | SE | t | $p_{bonf}$ |
|---|---|---|---|---|---|
| 1f | or | -0.989 | 0.534 | -1.853 | 0.198 |
| | ps | 3.017 | 0.534 | 5.654 | < .001 |
| or | ps | 4.006 | 0.534 | 7.507 | < .001 |

*Note.* P-value adjusted for comparing a family of 3
*Note.* Results are averaged over the levels of: cat

**Post Hoc Comparisons - manip ✻ cat**

| | | Mean Difference | SE | t | $p_{bonf}$ |
|---|---|---|---|---|---|
| 1f N | or N | -1.600 | 0.675 | -2.370 | 0.287 |
| | ps N | 5.069 | 0.675 | 7.509 | < .001 |
| | 1f T | 4.088 | 0.755 | 5.416 | < .001 |
| | or T | 3.710 | 0.755 | 4.916 | < .001 |
| | ps T | 5.053 | 0.755 | 6.696 | < .001 |
| or N | ps N | 6.669 | 0.675 | 9.879 | < .001 |
| | 1f T | 5.688 | 0.755 | 7.536 | < .001 |
| | or T | 5.310 | 0.755 | 7.036 | < .001 |
| | ps T | 6.654 | 0.755 | 8.816 | < .001 |
| ps N | 1f T | -0.981 | 0.755 | -1.300 | 1.000 |
| | or T | -1.359 | 0.755 | -1.801 | 1.000 |
| | ps T | -0.016 | 0.755 | -0.021 | 1.000 |
| 1f T | or T | -0.378 | 0.827 | -0.457 | 1.000 |
| | ps T | 0.965 | 0.827 | 1.168 | 1.000 |
| or T | ps T | 1.343 | 0.827 | 1.625 | 1.000 |

*Note.* P-value adjusted for comparing a family of 6

**Supplementary Table S3:** ANOVA for the effects of image category (non-texture: **N**, texture: **T**) and manipulation (**or**: original, **ps**: phase-scrambled, **1f**: amplitude-normalized) on the kurtosis of the log-Gabor filter population (**kurlog**) for the assumed viewing distance of 6 dva.

#### SUPPLEMENTARY TABLE S4

**ANOVA - actlog**

| Cases | Sum of Squares | df | Mean Square | F | p | η² |
|---|---|---|---|---|---|---|
| manip | 3.246 | 2 | 1.623 | 4.788 | 0.010 | 0.026 |
| cat | 57.123 | 1 | 57.123 | 168.537 | < .001 | 0.459 |
| manip ✻ cat | 15.320 | 2 | 7.660 | 22.601 | < .001 | 0.123 |
| Residuals | 48.806 | 144 | 0.339 | | | |

*Note.* Type III Sum of Squares

**Post Hoc Comparisons - manip**

| | | Mean Difference | SE | t | $p_{bonf}$ |
|---|---|---|---|---|---|
| 1f | or | 0.149 | 0.119 | 1.258 | 0.631 |
| | ps | -0.216 | 0.119 | -1.820 | 0.213 |
| or | ps | -0.366 | 0.119 | -3.078 | 0.007 |

*Note.* P-value adjusted for comparing a family of 3
*Note.* Results are averaged over the levels of: cat

**Post Hoc Comparisons - manip ✻ cat**

| | | Mean Difference | SE | t | $p_{bonf}$ |
|---|---|---|---|---|---|
| 1f N | or N | 0.914 | 0.150 | 6.083 | < .001 |
| | ps N | 0.366 | 0.150 | 2.435 | 0.242 |
| | 1f T | -0.362 | 0.168 | -2.151 | 0.497 |
| | or T | -0.977 | 0.168 | -5.813 | < .001 |
| | ps T | -1.160 | 0.168 | -6.902 | < .001 |
| or N | ps N | -0.548 | 0.150 | -3.649 | 0.006 |
| | 1f T | -1.276 | 0.168 | -7.592 | < .001 |
| | or T | -1.891 | 0.168 | -11.254 | < .001 |
| | ps T | -2.074 | 0.168 | -12.343 | < .001 |
| ps N | 1f T | -0.728 | 0.168 | -4.329 | < .001 |
| | or T | -1.343 | 0.168 | -7.991 | < .001 |
| | ps T | -1.526 | 0.168 | -9.080 | < .001 |
| 1f T | or T | -0.615 | 0.184 | -3.343 | 0.016 |
| | ps T | -0.798 | 0.184 | -4.337 | < .001 |
| or T | ps T | -0.183 | 0.184 | -0.994 | 1.000 |

*Note.* P-value adjusted for comparing a family of 6

**Supplementary Table S4:** ANOVA for the effects of image category (non-texture: **N**, texture: **T**) and manipulation (**or**: original, **ps**: phase-scrambled, **1f**: amplitude-normalized) on the kurtosis of the log-Gabor filter population (**actlog**) for the assumed viewing distance of 6 dva.

#### SUPPLEMENTARY TABLE S5

**ANOVA - kurlog**

| Cases | Sum of Squares | df | Mean Square | F | p | η² |
|---|---|---|---|---|---|---|
| cat | 507.322 | 1 | 507.322 | 44.418 | < .001 | 0.167 |
| manip | 617.855 | 2 | 308.928 | 27.048 | < .001 | 0.203 |
| cat ✻ manip | 273.412 | 2 | 136.706 | 11.969 | < .001 | 0.090 |
| Residuals | 1644.710 | 144 | 11.422 | | | |

*Note.* Type III Sum of Squares

**Post Hoc Comparisons - manip**

| | | Mean Difference | SE | t | $p_{bonf}$ |
|---|---|---|---|---|---|
| 1f | or | -1.327 | 0.690 | -1.924 | 0.169 |
| | ps | 3.578 | 0.690 | 5.186 | < .001 |
| or | ps | 4.905 | 0.690 | 7.110 | < .001 |

*Note.* P-value adjusted for comparing a family of 3
*Note.* Results are averaged over the levels of: cat

**Post Hoc Comparisons - cat ✻ manip**

| | | Mean Difference | SE | t | $p_{bonf}$ |
|---|---|---|---|---|---|
| N 1f | T 1f | 4.764 | 0.976 | 4.883 | < .001 |
| | N or | -2.199 | 0.873 | -2.521 | 0.192 |
| | T or | 4.309 | 0.976 | 4.417 | < .001 |
| | N ps | 5.965 | 0.873 | 6.836 | < .001 |
| | T ps | 5.954 | 0.976 | 6.103 | < .001 |
| T 1f | N or | -6.963 | 0.976 | -7.138 | < .001 |
| | T or | -0.455 | 1.069 | -0.425 | 1.000 |
| | N ps | 1.201 | 0.976 | 1.231 | 1.000 |
| | T ps | 1.190 | 1.069 | 1.114 | 1.000 |
| N or | T or | 6.509 | 0.976 | 6.672 | < .001 |
| | N ps | 8.165 | 0.873 | 9.357 | < .001 |
| | T ps | 8.154 | 0.976 | 8.357 | < .001 |
| T or | N ps | 1.656 | 0.976 | 1.697 | 1.000 |
| | T ps | 1.645 | 1.069 | 1.539 | 1.000 |
| N ps | T ps | -0.011 | 0.976 | -0.011 | 1.000 |

*Note.* P-value adjusted for comparing a family of 6

**Supplementary Table S5:** ANOVA for the effects of image category (non-texture: **N**, texture: **T**) and manipulation (**or**: original, **ps**: phase-scrambled, **1f**: amplitude-normalized) on the kurtosis of the log-Gabor filter population (**kurlog**) for the assumed viewing distance of 8 dva.

### SUPPLEMENTARY TABLE S6

**ANOVA - actlog**

| Cases | Sum of Squares | df | Mean Square | F | p | η² |
|---|---|---|---|---|---|---|
| cat | 80.589 | 1 | 80.589 | 235.610 | < .001 | 0.513 |
| manip | 3.732 | 2 | 1.866 | 5.456 | 0.005 | 0.024 |
| cat ✻ manip | 23.367 | 2 | 11.683 | 34.157 | < .001 | 0.149 |
| Residuals | 49.254 | 144 | 0.342 | | | |

*Note.* Type III Sum of Squares

**Post Hoc Comparisons - manip**

| | | Mean Difference | SE | t | $p_{bonf}$ |
|---|---|---|---|---|---|
| 1f | or | 0.206 | 0.119 | 1.726 | 0.259 |
| | ps | -0.188 | 0.119 | -1.576 | 0.352 |
| or | ps | -0.394 | 0.119 | -3.302 | 0.004 |

*Note.* P-value adjusted for comparing a family of 3
*Note.* Results are averaged over the levels of: cat

**Post Hoc Comparisons - cat ✻ manip**

| | | Mean Difference | SE | t | $p_{bonf}$ |
|---|---|---|---|---|---|
| N 1f | T 1f | -0.375 | 0.169 | -2.221 | 0.419 |
| | N or | 1.135 | 0.151 | 7.514 | < .001 |
| | T or | -1.097 | 0.169 | -6.500 | < .001 |
| | N ps | 0.565 | 0.151 | 3.743 | 0.004 |
| | T ps | -1.316 | 0.169 | -7.797 | < .001 |
| T 1f | N or | 1.510 | 0.169 | 8.942 | < .001 |
| | T or | -0.722 | 0.185 | -3.906 | 0.002 |
| | N ps | 0.940 | 0.169 | 5.569 | < .001 |
| | T ps | -0.941 | 0.185 | -5.090 | < .001 |
| N or | T or | -2.232 | 0.169 | -13.221 | < .001 |
| | N ps | -0.569 | 0.151 | -3.771 | 0.004 |
| | T ps | -2.451 | 0.169 | -14.518 | < .001 |
| T or | N ps | 1.663 | 0.169 | 9.848 | < .001 |
| | T ps | -0.219 | 0.185 | -1.184 | 1.000 |
| N ps | T ps | -1.882 | 0.169 | -11.145 | < .001 |

*Note.* P-value adjusted for comparing a family of 6

**Supplementary Table S6:** ANOVA for the effects of image category (non-texture: **N**, texture: **T**) and manipulation (**or**: original, **ps**: phase-scrambled, **1f**: amplitude-normalized) on the activity of the log-Gabor filter population (**actlog**) for the assumed viewing distance of 8 dva.

**SUPPLEMENTARY TABLE S7**

| | **mSF** | **act-log** | **kur-log** |
|---|---|---|---|
| **mSF** | - | - | - |
| **act-log** | 0.898*** | - | - |
| **kur-log** | -0.541*** | -0.683*** | - |

* = p < 0.05, ** = p < 0.01, *** = p < 0.001

**Supplementary Table S7:** Significant correlations between image statistics for the original images (**or**) for 4 dva assumption of image size.

**SUPPLEMENTARY TABLE S8**

| | **mSF** | **act-log** | **kur-log** |
|---|---|---|---|
| **mSF** | - | - | - |
| **act-log** | 0.956*** | - | - |
| **kur-log** | -0.637*** | -0.707*** | - |

* = p < 0.05, ** = p < 0.01, *** = p < 0.001

**Supplementary Table S8:** Significant correlations between image statistics for the original images (**or**) for 8 dva assumption of image size.

### SUPPLEMENTARY TABLE S9

*ANOVA - comfort*

| Cases | Sum of Squares | df | Mean Square | F | p | $\eta^2$ |
|---|---|---|---|---|---|---|
| cat | 27.59 | 1 | 27.59 | 74.65 | < .001 | 0.150 |
| manip | 91.15 | 2 | 45.58 | 123.3 | < .001 | 0.497 |
| cat ✻ manip | 11.46 | 2 | 5.732 | 15.51 | < .001 | 0.062 |
| Residuals | 53.23 | 144 | 0.370 | | | |

Note. Type III Sum of Squares

*Descriptives - comfort*

| cat | manip | N | Mean | SD | SE | Coefficient of variation |
|---|---|---|---|---|---|---|
| N | 1f | 30 | 1.149 | 0.580 | 0.106 | 0.505 |
| | or | 30 | 2.457 | 0.676 | 0.123 | 0.275 |
| | ps | 30 | -0.180 | 0.176 | 0.032 | -0.978 |
| T | 1f | 20 | 0.202 | 0.528 | 0.118 | 2.608 |
| | or | 20 | 0.928 | 1.101 | 0.246 | 1.186 |
| | ps | 20 | -0.331 | 0.231 | 0.052 | -0.697 |

*Post Hoc Comparisons - cat ✻ manip*

| | | Mean Difference | SE | df | t | pBonf |
|---|---|---|---|---|---|---|
| N 1f | T 1f | 0.946 | 0.176 | 144 | 5.393 | < .001 |
| | N or | -1.308 | 0.157 | 144 | -8.333 | < .001 |
| | T or | 0.220 | 0.176 | 144 | 1.255 | 1.000 |
| | N ps | 1.329 | 0.157 | 144 | 8.463 | < .001 |
| | T ps | 1.480 | 0.176 | 144 | 8.433 | < .001 |
| T 1f | N or | -2.255 | 0.176 | 144 | -12.85 | < .001 |
| | T or | -0.726 | 0.192 | 144 | -3.777 | .003 |
| | N ps | 0.382 | 0.176 | 144 | 2.177 | .467 |
| | T ps | 0.534 | 0.192 | 144 | 2.776 | .094 |
| N or | T or | 1.528 | 0.176 | 144 | 8.708 | < .001 |
| | N ps | 2.637 | 0.157 | 144 | 16.80 | < .001 |
| | T ps | 2.788 | 0.176 | 144 | 15.89 | < .001 |
| T or | N ps | 1.108 | 0.176 | 144 | 6.314 | < .001 |
| | T ps | 1.260 | 0.192 | 144 | 6.553 | < .001 |
| N ps | T ps | 0.152 | 0.176 | 144 | 0.864 | 1.000 |

Note. P-value adjusted for comparing a family of 15 estimates.

*Post Hoc Comparisons - manip*

| | | Mean Difference | SE | df | t | pBonf |
|---|---|---|---|---|---|---|
| 1f | or | -1.017 | 0.124 | 144 | -8.195 | < .001 |
| | ps | 0.931 | 0.124 | 144 | 7.503 | < .001 |
| or | ps | 1.948 | 0.124 | 144 | 15.70 | < .001 |

Note. P-value adjusted for comparing a family of 3 estimates.
Note. Results are averaged over the levels of: cat

**Supplementary Table S9:** ANOVA for comfort ratings with factors of category (**T**: texture, **N**: non-textures) and manipulation (**or**: original, **ps**: phase-scrambeld , **1f**: amplitude-normalized) with all post-hoc comparisons.

### SUPPLEMENTARY TABLE S10

*Model Summary - comfort*

| Model | R | $R^2$ | Adjusted $R^2$ | RMSE | $R^2$ Change | df1 | df2 | p |
|---|---|---|---|---|---|---|---|---|
| $M_0$ | 0.000 | 0.000 | 0.000 | 1.146 | 0.000 | 0 | 49 | |
| $M_1$ | 0.346 | 0.120 | 0.102 | 1.086 | 0.120 | 1 | 48 | .014 |

Note. $M_1$ includes amp1oct3

*ANOVA*

| Model | | Sum of Squares | df | Mean Square | F | p |
|---|---|---|---|---|---|---|
| $M_1$ | Regression | 7.710 | 1 | 7.710 | 6.538 | .014 |
| | Residual | 56.60 | 48 | 1.179 | | |
| | Total | 64.31 | 49 | | | |

Note. $M_1$ includes amp1oct3
Note. The intercept model is omitted, as no meaningful information can be shown.

*Coefficients*

| Model | | Unstandardized | Standard Error | Standardized | t | p |
|---|---|---|---|---|---|---|
| $M_0$ | (Intercept) | 1.845 | 0.162 | | 11.39 | < .001 |
| $M_1$ | (Intercept) | 2.756 | 0.388 | | 7.107 | < .001 |
| | amp1oct3 | -0.866 | 0.339 | -0.346 | -2.557 | .014 |

**Supplementary Table S10:** Linear regression for viewing comfort for all original (**or**) images (both categories) as a function of mid-SF ratio (**amp1oct3**) for 4 dva viewing assumption.

### SUPPLEMENTARY TABLE S11

*Model Summary - comfort*

| Model | R | $R^2$ | Adjusted $R^2$ | RMSE | $R^2$ Change | df1 | df2 | p |
|---|---|---|---|---|---|---|---|---|
| $M_0$ | 0.000 | 0.000 | 0.000 | 1.146 | 0.000 | 0 | 49 | |
| $M_1$ | 0.525 | 0.276 | 0.261 | 0.985 | 0.276 | 1 | 48 | < .001 |

Note. $M_1$ includes amp1oct3

*ANOVA*

| Model | | Sum of Squares | df | Mean Square | F | p |
|---|---|---|---|---|---|---|
| $M_1$ | Regression | 17.73 | 1 | 17.73 | 18.27 | < .001 |
| | Residual | 46.58 | 48 | 0.970 | | |
| | Total | 64.31 | 49 | | | |

Note. $M_1$ includes amp1oct3
Note. The intercept model is omitted, as no meaningful information can be shown.

*Coefficients*

| Model | | Unstandardized | Standard Error | Standardized | t | p |
|---|---|---|---|---|---|---|
| $M_0$ | (Intercept) | 1.845 | 0.162 | | 11.39 | < .001 |
| $M_1$ | (Intercept) | 2.910 | 0.285 | | 10.20 | < .001 |
| | amp1oct3 | -0.990 | 0.232 | -0.525 | -4.274 | < .001 |

**Supplementary Table S11:** Linear regression for viewing comfort for all original (**or**) images (both categories) as a function of mid-SF ratio (**amp1oct3**) for 6 dva viewing assumption.

#### SUPPLEMENTARY TABLE S12

*Model Summary - comfort*

| Model | R | $R^2$ | Adjusted $R^2$ | RMSE | $R^2$ Change | df1 | df2 | p |
|---|---|---|---|---|---|---|---|---|
| $M_0$ | 0.000 | 0.000 | 0.000 | 1.146 | 0.000 | 0 | 49 | |
| $M_1$ | 0.573 | 0.328 | 0.314 | 0.949 | 0.328 | 1 | 48 | < .001 |

Note. $M_1$ includes amp1oct3

*ANOVA*

| Model | | Sum of Squares | df | Mean Square | F | p |
|---|---|---|---|---|---|---|
| $M_1$ | Regression | 21.09 | 1 | 21.09 | 23.41 | < .001 |
| | Residual | 43.23 | 48 | 0.901 | | |
| | Total | 64.31 | 49 | | | |

Note. $M_1$ includes amp1oct3
Note. The intercept model is omitted, as no meaningful information can be shown.

*Coefficients*

| Model | | Unstandardized | Standard Error | Standardized | t | p |
|---|---|---|---|---|---|---|
| $M_0$ | (Intercept) | 1.845 | 0.162 | | 11.39 | < .001 |
| $M_1$ | (Intercept) | 2.846 | 0.247 | | 11.54 | < .001 |
| | amp1oct3 | -0.930 | 0.192 | -0.573 | -4.839 | < .001 |

**Supplementary Table S12:** Linear regression for viewing comfort for all original (**or**) images (both categories) as a function of mid-SF ratio (**amp1oct3**) for 8 dva viewing assumption.

**SUPPLEMENTARY TABLE S13**

**Model Summary - comfort**

| Model | R | $R^2$ | Adjusted $R^2$ | RMSE |
|---|---|---|---|---|
| $H_0$ | 0.000 | 0.000 | 0.000 | 1.146 |
| $H_1$ | 0.508 | 0.258 | 0.226 | 1.008 |

**ANOVA**

| Model | | Sum of Squares | df | Mean Square | F | p |
|---|---|---|---|---|---|---|
| $H_1$ | Regression | 16.575 | 2 | 8.287 | 8.160 | < .001 |
| | Residual | 47.736 | 47 | 1.016 | | |
| | Total | 64.310 | 49 | | | |

*Note.* The intercept model is omitted, as no meaningful information can be shown.

**Coefficients**

| Model | | Unstandardized | Standard Error | Standardized | t | p |
|---|---|---|---|---|---|---|
| $H_0$ | (Intercept) | 1.845 | 0.162 | | 11.391 | < .001 |
| $H_1$ | (Intercept) | 2.402 | 1.188 | | 2.021 | 0.049 |
| | kurlog | 0.116 | 0.068 | 0.292 | 1.699 | 0.096 |
| | actlog | -0.325 | 0.214 | -0.261 | -1.519 | 0.135 |

**Supplementary Table S13:** Linear regression for viewing comfort for all original (**or**) images (both categories) as a function of **kur-log** and **act-log** for 4 dva viewing assumption.

**SUPPLEMENTARY TABLE S14**

**Model Summary - comfort**

| Model | R | $R^2$ | Adjusted $R^2$ | RMSE |
|---|---|---|---|---|
| $H_0$ | 0.000 | 0.000 | 0.000 | 1.146 |
| $H_1$ | 0.577 | 0.333 | 0.304 | 0.956 |

**ANOVA**

| Model | | Sum of Squares | df | Mean Square | F | p |
|---|---|---|---|---|---|---|
| $H_1$ | Regression | 21.395 | 2 | 10.697 | 11.716 | < .001 |
| | Residual | 42.916 | 47 | 0.913 | | |
| | Total | 64.310 | 49 | | | |

*Note.* The intercept model is omitted, as no meaningful information can be shown.

**Coefficients**

| Model | | Unstandardized | Standard Error | Standardized | t | p |
|---|---|---|---|---|---|---|
| $H_0$ | (Intercept) | 1.845 | 0.162 | | 11.391 | < .001 |
| $H_1$ | (Intercept) | 3.602 | 0.931 | | 3.867 | < .001 |
| | kurlog | 0.022 | 0.046 | 0.081 | 0.471 | 0.640 |
| | actlog | -0.504 | 0.168 | -0.516 | -3.000 | 0.004 |

**Supplementary Table S14:** Linear regression for viewing comfort for all original (**or**) images (both categories) as a function of **kur-log** and **act-log** for 6 dva viewing assumption.

**SUPPLEMENTARY TABLE S15**

**Model Summary - comfort**

| Model | R | $R^2$ | Adjusted $R^2$ | RMSE |
|---|---|---|---|---|
| $H_0$ | 0.000 | 0.000 | 0.000 | 1.146 |
| $H_1$ | 0.611 | 0.373 | 0.346 | 0.926 |

**ANOVA**

| Model | | Sum of Squares | df | Mean Square | F | p |
|---|---|---|---|---|---|---|
| $H_1$ | Regression | 23.984 | 2 | 11.992 | 13.977 | < .001 |
| | Residual | 40.326 | 47 | 0.858 | | |
| | Total | 64.310 | 49 | | | |

*Note.* The intercept model is omitted, as no meaningful information can be shown.

**Coefficients**

| Model | | Unstandardized | Standard Error | Standardized | t | p |
|---|---|---|---|---|---|---|
| $H_0$ | (Intercept) | 1.845 | 0.162 | | 11.391 | < .001 |
| $H_1$ | (Intercept) | 3.780 | 0.767 | | 4.925 | < .001 |
| | actlog | -0.528 | 0.142 | -0.606 | -3.714 | < .001 |
| | kurlog | 0.001 | 0.034 | 0.006 | 0.038 | 0.970 |

**Supplementary Table S15:** Linear regression for viewing comfort for all original (**or**) images (both categories) as a function of **kur-log** and **act-log** for 8 dva viewing assumption.

#### SUPPLEMENTARY TABLE S16

Indirect and Total Effects

| | | | | 95% C.I. (a) | | | | |
|---|---|---|---|---|---|---|---|---|
| Type | Effect | Estimate | SE | Lower | Upper | β | z | p |
| Indirect | amp1oct3 ⇒ actlog ⇒ comfort | -1.733 | 0.642 | -2.991 | -0.475 | -0.692 | -2.70 | 0.007 |
| Component | amp1oct3 ⇒ actlog | 1.805 | 0.125 | 1.559 | 2.051 | 0.898 | 14.40 | < .001 |
| | actlog ⇒ comfort | -0.960 | 0.349 | -1.645 | -0.276 | -0.771 | -2.75 | 0.006 |
| Direct | amp1oct3 ⇒ comfort | 0.866 | 0.702 | -0.510 | 2.243 | 0.346 | 1.23 | 0.217 |
| Total | amp1oct3 ⇒ comfort | -0.866 | 0.335 | -1.524 | -0.209 | -0.346 | -2.58 | 0.010 |

Note. Confidence intervals computed with method: Standard (Delta method)

Note. Betas are completely standardized effect sizes

**Supplementary Table S16:** Mediation analysis for 4 dva viewing distance assumption.

#### SUPPLEMENTARY TABLE S17

Indirect and Total Effects

| Type | Effect | Estimate | SE | 95% C.I. (a) Lower | 95% C.I. (a) Upper | β | z | p |
|---|---|---|---|---|---|---|---|---|
| Indirect | amp1oct3 ⇒ actlog ⇒ comfort | -1.225 | 0.6035 | -2.41 | -0.0421 | -0.650 | -2.030 | 0.042 |
| Component | amp1oct3 ⇒ actlog | 1.811 | 0.0930 | 1.63 | 1.9937 | 0.940 | 19.471 | < .001 |
| | actlog ⇒ comfort | -0.676 | 0.3314 | -1.33 | -0.0268 | -0.691 | -2.041 | 0.041 |
| Direct | amp1oct3 ⇒ comfort | 0.235 | 0.6386 | -1.02 | 1.4869 | 0.125 | 0.368 | 0.713 |
| Total | amp1oct3 ⇒ comfort | -0.990 | 0.2292 | -1.44 | -0.5404 | -0.525 | -4.318 | < .001 |

Note. Confidence intervals computed with method: Standard (Delta method)

Note. Betas are completely standardized effect sizes

**Supplementary Table S17:** Mediation analysis for 6 dva viewing distance assumption.

### SUPPLEMENTARY TABLE S18

Indirect and Total Effects

| Type | Effect | Estimate | SE | 95% C.I. (a) Lower | 95% C.I. (a) Upper | β | z | p |
|---|---|---|---|---|---|---|---|---|
| Indirect | amp1oct3 ⇒ actlog ⇒ comfort | -1.145 | 0.5955 | -2.31 | 0.0223 | -0.705 | -1.922 | 0.055 |
| Component | amp1oct3 ⇒ actlog | 1.782 | 0.0772 | 1.63 | 1.9336 | 0.956 | 23.098 | < .001 |
| | actlog ⇒ comfort | -0.642 | 0.3329 | -1.29 | 0.0103 | -0.737 | -1.929 | 0.054 |
| Direct | amp1oct3 ⇒ comfort | 0.215 | 0.6206 | -1.00 | 1.4312 | 0.132 | 0.346 | 0.729 |
| Total | amp1oct3 ⇒ comfort | -0.930 | 0.1902 | -1.30 | -0.5571 | -0.573 | -4.889 | < .001 |

Note. Confidence intervals computed with method: Standard (Delta method)

Note. Betas are completely standardized effect sizes

**Supplementary Table S18:** Mediation analysis for 8 dva viewing distance assumption.

#### SUPPLEMENTARY TABLE S19

Model Fit Measures

| Model | R | $R^2$ |
|---|---|---|
| 1 | 0.662 | 0.438 |

Model Coefficients - comfort

| Predictor | Estimate | SE | t | p |
|---|---|---|---|---|
| Intercept [a] | 2.6950 | 0.641 | 4.206 | < .001 |
| actlog | -0.0687 | 0.179 | -0.384 | 0.703 |
| cat: | | | | |
| T – N | -1.4454 | 0.333 | -4.341 | < .001 |

[a] Represents reference level

**Supplementary Table S19:** Linear regression with factors of category (**cat**) and co-variate of **act-log** for dva = 4.

#### SUPPLEMENTARY TABLE S20

Model Fit Measures

| Model | R | $R^2$ |
|---|---|---|
| 1 | 0.665 | 0.442 |

Model Coefficients - comfort

| Predictor | Estimate | SE | t | p |
|---|---|---|---|---|
| Intercept [a] | 2.840 | 0.564 | 5.039 | < .001 |
| actlog | -0.126 | 0.177 | -0.709 | 0.482 |
| cat: | | | | |
| T – N | -1.291 | 0.420 | -3.076 | 0.003 |

[a] Represents reference level

**Supplementary Table S20:** Linear regression with factors of category (**cat**) and co-variate of **act-log** for dva = 6.

#### SUPPLEMENTARY TABLE S21

Model Fit Measures

| Model | R | $R^2$ |
|---|---|---|
| 1 | 0.668 | 0.447 |

Model Coefficients - comfort

| Predictor | Estimate | SE | t | p |
|---|---|---|---|---|
| Intercept [a] | 2.920 | 0.511 | 5.709 | < .001 |
| actlog | -0.166 | 0.174 | -0.952 | 0.346 |
| cat: | | | | |
| T – N | -1.158 | 0.463 | -2.501 | 0.016 |

[a] Represents reference level

**Supplementary Table S21:** Linear regression with factors of category (**cat**) and co-variate of **act-log** for dva = 8.

#### SUPPLEMENTARY TABLE S22

Model Info

| Info | |
|---|---|
| Estimate | Linear mixed model fit by REML |
| Call | comfort ~ 1 + manip + cat + actlog + manip:cat+( 1 \| image )+( 1 \| Subject ) |
| AIC | 81447.032 |
| BIC | 81552.603 |
| LogLikel. | -40726.834 |
| R-squared Marginal | 0.136 |
| R-squared Conditional | 0.531 |
| Converged | yes |
| Optimizer | bobyqa |

Fixed Effect Omnibus tests

| | F | Num df | Den df | p |
|---|---|---|---|---|
| manip | 1442.6 | 2 | 19248.5 | < .001 |
| cat | 32.0 | 1 | 53.6 | < .001 |
| actlog | 19.8 | 1 | 4102.9 | < .001 |
| manip ✻ cat | 204.3 | 2 | 18537.0 | < .001 |

Note. Satterthwaite method for degrees of freedom

**Supplementary Table S22:** Linear mixed effects model including **act-log** (1-way model) for 4 dva viewing distance assumption.

#### SUPPLEMENTARY TABLE S23

Model Info

| Info | |
|---|---|
| Estimate | Linear mixed model fit by REML |
| Call | comfort ~ 1 + cat + manip + actlog + cat:manip+( 1 \| image )+( 1 \| Subject ) |
| AIC | 81436.100 |
| BIC | 81541.620 |
| LogLikel. | -40721.343 |
| R-squared Marginal | 0.137 |
| R-squared Conditional | 0.532 |
| Converged | yes |
| Optimizer | bobyqa |

Fixed Effect Omnibus tests

| | F | Num df | Den df | p |
|---|---|---|---|---|
| cat | 18.7 | 1 | 63.6 | < .001 |
| manip | 1326.6 | 2 | 18658.6 | < .001 |
| actlog | 30.8 | 1 | 3583.5 | < .001 |
| cat ✻ manip | 217.4 | 2 | 12937.3 | < .001 |

Note. Satterthwaite method for degrees of freedom

**Supplementary Table S23:** Linear mixed effects model including **act-log** (1-way model) for 6 dva viewing distance assumption.

#### SUPPLEMENTARY TABLE S24

Model Info

| Info | |
|---|---|
| Estimate | Linear mixed model fit by REML |
| Call | comfort ~ 1 + cat + manip + actlog + cat:manip+( 1 \| image )+( 1 \| Subject ) |
| AIC | 81430.438 |
| BIC | 81535.954 |
| LogLikel. | -40718.510 |
| R-squared Marginal | 0.137 |
| R-squared Conditional | 0.532 |
| Converged | yes |
| Optimizer | bobyqa |

Fixed Effect Omnibus tests

| | F | Num df | Den df | p |
|---|---|---|---|---|
| cat | 12.8 | 1 | 71.1 | < .001 |
| manip | 1261.1 | 2 | 18131.3 | < .001 |
| actlog | 36.4 | 1 | 3267.5 | < .001 |
| cat ✻ manip | 224.6 | 2 | 10375.4 | < .001 |

Note. Satterthwaite method for degrees of freedom

**Supplementary Table S24:** Linear mixed effects model including **act-log** (1-way model) for 8 dva viewing distance assumption.

### SUPPLEMENTARY TABLE S25

Model Info

| Info | |
|---|---|
| Estimate | Linear mixed model fit by REML |
| Call | comfort ~ 1 + manip + cat + actlog + manip:cat + cat:actlog + manip:actlog+( 1 \| image )+( 1 \| Subject ) |
| AIC | 81442.053 |
| BIC | 81579.933 |
| LogLikel. | -40725.659 |
| R-squared Marginal | 0.136 |
| R-squared Conditional | 0.532 |
| Converged | yes |
| Optimizer | bobyqa |

Fixed Effect Omnibus tests

| | F | Num df | Den df | p |
|---|---|---|---|---|
| manip | 1395.46 | 2 | 16848.6 | < .001 |
| cat | 23.80 | 1 | 59.9 | < .001 |
| actlog | 15.31 | 1 | 5603.2 | < .001 |
| manip ✻ cat | 141.05 | 2 | 18667.8 | < .001 |
| cat ✻ actlog | 1.52 | 1 | 4522.5 | 0.217 |
| manip ✻ actlog | 4.73 | 2 | 15553.6 | 0.009 |

Note. Satterthwaite method for degrees of freedom

**Supplementary Table S25:** Linear mixed effects model including **act-log** (2-way model) for 4 dva viewing distance assumption.

#### SUPPLEMENTARY TABLE S26

Model Info

| Info | |
|---|---|
| Estimate | Linear mixed model fit by REML |
| Call | comfort ~ 1 + cat + manip + actlog + cat:manip + cat:actlog + manip:actlog+( 1 \| image )+( 1 \| Subject ) |
| AIC | 81424.587 |
| BIC | 81562.520 |
| LogLikel. | -40716.953 |
| R-squared Marginal | 0.138 |
| R-squared Conditional | 0.533 |
| Converged | yes |
| Optimizer | bobyqa |

Fixed Effect Omnibus tests

| | F | Num df | Den df | p |
|---|---|---|---|---|
| cat | 7.70 | 1 | 79.4 | 0.007 |
| manip | 1236.06 | 2 | 11490.3 | < .001 |
| actlog | 34.17 | 1 | 3761.9 | < .001 |
| cat ✻ manip | 95.23 | 2 | 17633.1 | < .001 |
| cat ✻ actlog | 3.51 | 1 | 3975.9 | 0.061 |
| manip ✻ actlog | 7.22 | 2 | 14205.4 | < .001 |

Note. Satterthwaite method for degrees of freedom

**Supplementary Table S26:** Linear mixed effects model including **act-log** (2-way model) for 6 dva viewing distance assumption.

### SUPPLEMENTARY TABLE S27

Model Info

| Info | |
|---|---|
| Estimate | Linear mixed model fit by REML |
| Call | comfort ~ 1 + cat + manip + actlog + cat:manip + cat:actlog + manip:actlog+( 1 \| image )+( 1 \| Subject ) |
| AIC | 81415.701 |
| BIC | 81553.818 |
| LogLikel. | -40712.602 |
| R-squared Marginal | 0.139 |
| R-squared Conditional | 0.533 |
| Converged | yes |
| Optimizer | bobyqa |

Fixed Effect Omnibus tests

| | F | Num df | Den df | p |
|---|---|---|---|---|
| cat | 3.17 | 1 | 90.6 | 0.078 |
| manip | 1161.77 | 2 | 9383.5 | < .001 |
| actlog | 46.42 | 1 | 2957.0 | < .001 |
| cat ✻ manip | 75.20 | 2 | 16232.9 | < .001 |
| cat ✻ actlog | 3.07 | 1 | 3539.8 | 0.080 |
| manip ✻ actlog | 9.09 | 2 | 13162.3 | < .001 |

Note. Satterthwaite method for degrees of freedom

**Supplementary Table S27:** Linear mixed effects model including **act-log** (2-way model) for 8 dva viewing distance assumption.

**SUPPLEMENTARY FIGURE S1**

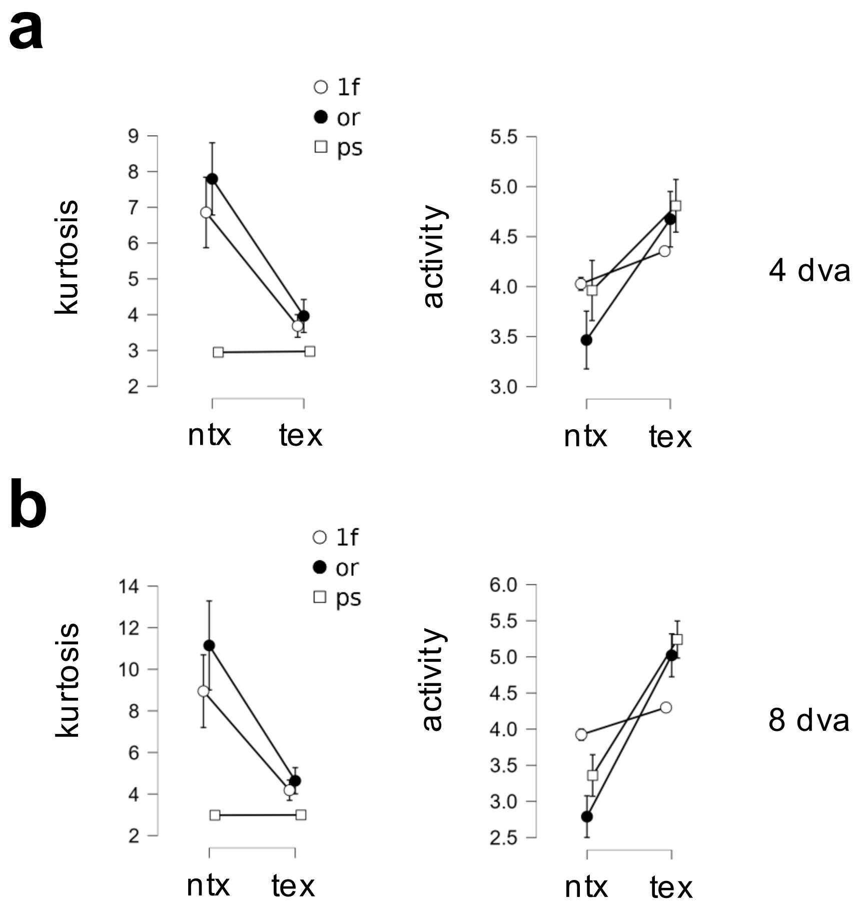


**Supplementary Fig. S1:** Same as **Fig. 4a** in main text, but for 4, 8 dva viewing distance assumptions.

#### SUPPLEMENTARY FIGURE S2

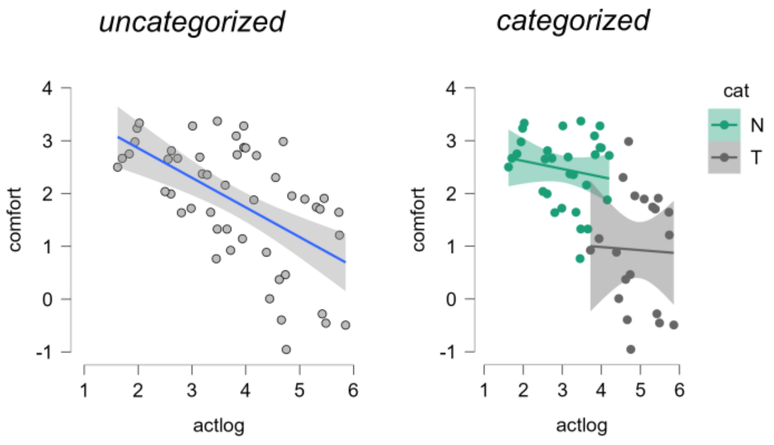


**Supplementary Fig. S2:** Relationship between activity of log-Gabor filter population activity and viewing comfort averaged across observers for original images (6 dva assumption), when image category is not a factor (left columns) or is included as a factor (right columns).
